## Supporting Information for "Deciphering regulatory architectures from synthetic single-cell expression patterns"

[1]Division of Biology and Biological Engineering, California Institute of Technology, Pasadena, CA; [2]Biophysics Graduate Group, University of California, Berkeley, CA; [3]Department of Physics, University of California, Berkeley, CA; [4]Department of Molecular and Cell Biology, University of California, Berkeley, CA; [5]Institute for Quantitative Biosciences-QB3, University of California, Berkeley, CA; [6]Department of Physics, California Institute of Technology, Pasadena, CA

#### S1 Appendix Models of the probabilities of microscopic states during transcription

For the purposes of this paper, the first step in the modeling of transcription is the identification of some set of microscopic states that represent the entire set of states available to the promoter dictating our gene of interest. Already, such models represent a coarse-graining of the complex microscopic reality of DNA and its attendant proteins within a cell. For example, DNA conformation at all scales (chromatin structure, supercoiling, etc.) is not included in the most naive application of these models. This kind of coarse-graining is consistent with the depiction of regulatory architectures as a region of DNA with some collection of binding sites and their allied transcription factors such as seen in RegulonDB or EcoCyc and represented in Fig 3 in the paper. Though the assumption of discrete states itself deserves further scrutiny, in this paper we accept these assumptions about a set of discrete states with no further discussion. Once we accept this constellation of microscopic states, many models focus on the steady-state probabilities of those states. Of course, the full dynamical trajectories of these different microstates are of great interest and one approach to computing those dynamics is the use of coupled chemical master equations. With the advent of experimental approaches to measuring the *dynamics* of transcription, the analysis of gene expression dynamics requires something more than is offered by the steady-state probabilities considered here.

In general, the steady-state probabilities of the different microstates can be written in the form

$$p_i(x_1, x_2, \cdots x_n) = \frac{P_i(x_1, x_2, \cdots x_n)}{Q(x_1, x_2, \cdots x_n)}, \tag{S1}$$

where here we examine the probability of the $i^{th}$ state as a rational function (i.e. a ratio of two polynomials) where $x_j$ is the concentration of the $j^{th}$ transcription factor (or RNA polymerase) and with $Q(x_1, x_2, \cdots x_n)$ serving as a kind of generalized partition function gotten by summing over all states (or paths). In some instances, it is justified to specialize Eq S1 to the form

$$p_i(x) = \frac{P_i(x)}{Q(x)}, \tag{S2}$$

where for example our regulatory architecture of interest features only a single repressor or activator whose concentration is measured by $x$. If we think of the huge topic of input-output functions in biology, then $P_i(x)/Q(x)$ includes a representation of leakiness (the amount of output $p_i(x)$ even in the absence of input, $x = 0$), dynamic range, $EC_{50}$ (the concentration at which the output reaches half its maximum) and the sensitivity as measured by the slope of the input-output curve (usually in logarithmic variables) at the midpoint [1].

In the present paper, we exploit two different ways of thinking about these rational functions, one of which is a subset of the other. In particular, the most general models we consider provide the steady-state probabilities using the tools of graph theory and make no assumption of a quasi-equilibrium [2, 3]. Indeed, such models are a convenient platform for exploring the precise consequences of breaking detailed balance associated with some kinetic step connecting two different microstates [4]. Within the study of transcriptional regulation, the more common class of useful models is sometimes referred to as "thermodynamic models," which have a deep and interesting history in the context of both test-tube biochemistry and the study of signaling, regulation and physiology within living organisms. Such models have played an important role as a conceptual framework for more than a century and a convenient and inspiring point of departure is the work of Archibald V Hill after whom the famed Hill function

$$p_{\text{bound}}(x) = \frac{\left(\frac{x}{K}\right)^n}{1+\left(\frac{x}{K}\right)^n} \tag{S3}$$

is named. In this case, $x$ is the concentration of some ligand and $K$ is an effective dissociation constant. As Hill himself tells us, this functional form was hypothesized to describe the occupancy of hemoglobin by oxygen (the example he used, though it applies and has been applied much more broadly). Already more than a century ago, Hill argued of the function that now bears his name: "My object was rather to see whether an equation *of this type* can satisfy all the observations, than to base any direct physical meaning on $n$ and $K$." [5] He goes further in his 1913 paper noting "In point of fact $n$ does not turn out to be a whole number, but this is due simply to the fact that aggregation is not into one particular type of molecule, but rather into a whole series of different molecules: so that equation (1) is a rough mathematical expression for the sum of several similar quantities with $n$ equal to 1, 2, 3, 4 and possibly higher integers." [6]

In the subsequent decades, the equilibrium analysis of biochemical interactions became increasingly sophisticated with Pauling formulating a model that goes far beyond the Hill function by computing the average number of oxygen molecules bound to hemoglobin in the form

$$\langle N_{\text{bound}} \rangle = \frac{4x + 12x^2 j + 12x^3 j^3 + 4x^4 j^6}{1 + 4x + 6x^2 j + 4x^3 j^3 + x^4 j^6}, \tag{S4}$$

where we adopt the simplifying notation $x = [O_2]$ [7, 8]. The parameter $j$ is an interaction energy that imposes cooperativity in the sense that once one $O_2$ molecule is bound, the binding of the next one is easier. The equilibrium model of Adair went even further [9], positing that the average number of oxygen molecules bound to hemoglobin is given by

$$\langle N_{\text{bound}} \rangle = \frac{4x + 12x^2 j + 12x^3 j^3 k + 4x^4 j^6 k^4 l}{1 + 4x + 6x^2 j + 4x^3 j^3 k + x^4 j^6 k^4 l}, \tag{S5}$$

where now the parameter $k$ captures 3-body interactions between $O_2$ molecules and the parameter $l$ captures 4-body interactions [7]. For more than a century, equilibrium thinking has suffused the study of biochemical reactions, even when promoted from the sterile setting of test-tube biochemistry to the messy world of hemoglobin molecules enclosed within red blood cells that are themselves cycling rapidly through the circulatory systems of animals ranging from high-flying birds such as the bar-headed geese to elite divers such as blue whales.

It was not a huge leap to go from the idea of examples such as oxygen binding to hemoglobin (a special case of receptor-ligand binding) to the idea of DNA itself as the receptor and various proteins such as transcription factors and RNA polymerase as the "ligand." Classic work from Ackers and Shea [10, 11] formalized the kind of "regulated recruitment" thinking that had been diligently pursued by Ptashne and coworkers [12], turning it into a formal mathematical structure for evaluating the state probabilities for various occupancies of a promoter of interest. More recently, these ideas have been developed deeply by a battery of researchers several examples of which are given here [13, 14, 15]. Similarly, right from the get-go, the study of gene regulation made it clear that the phenomenon of induction (i.e. the use of effector molecules to tune the state of expression) would require a quantitative description, and beautiful work in the 1960s articulated a wide range of different equilibrium allosteric models such as the MWC model [16], the KNF model [17] and the generalization of these ideas in the model of Manfred Eigen [18]. All of these models, even if no one explicitly says so, are "thermodynamic models" in that they provide a systematic protocol for using statistical mechanics to find the probabilities of *all* of the allowed states. Note that these different models differ not in whether they are quasi-equilibrium or not, but rather in which states they permit.

The vast majority of approaches to generating a specific functional form for state probabilities in transcription like those given in Eq S1 are either: (a) some version of a thermodynamic model which appeals in one way or another to states and Boltzmann weights, (b) phenomenological guesses in which some convenient functional form is adopted (usually a Hill function) and more rarely, (c) using the tools of non-equilibrium physics and graph theory, a version of $p_i$ is adopted that reflects broken detailed balance. Approaches (a) and (c) both require some sort of mechanistic commitment about the classes of states that the system can adopt, and we note that (a) is a special case of (c). In the case of thermodynamic models, these polynomials have a very special form dictated by the Boltzmann weights of the different states of binding between transcription factors and their target DNA.

Part of the reason for the importance of this appendix is because there are so many different opinions in play on the subject of thermodynamic models in the biological setting in the literature. Part of the passion associated with the subject is that some authors are explicit in naming their work as thermodynamic or equilibrium models and others are not. Some fragment of the scientific population has an intrinsic belief that because "living organisms are out of equilibrium," thermodynamic or equilibrium models have no place. We think this extreme view can be replaced by a more nuanced perspective given that even within the fiery interior of a star, equilibrium ideas are used routinely and successfully (see the Saha equation). Rather, we are going to highlight several alternative and more nuanced pictures including: (i) justification based upon separation of time scales, (ii) a null model which serves as a first and simplest regulatory hypothesis and (iii) phenomenology.

First, we consider the status of thermodynamic models as a null model for the steady-state probabilities of biochemical phenomena with special emphasis on the application to transcription. Then, we subject such models to the harshest scrutiny: what is their track record as a conceptual framework for thinking about experimental data, and what is the nature of their shortcomings? Here we use the title "thermodynamic models" as a shorthand to refer to *all* models of occupancy of transcription factors, nucleosomes and polymerases in which the state probabilities are obtained from the Boltzmann distribution, or some phenomenological approximation (i.e. Hill function) to the equilibrium state probabilities. These models are ubiquitous not only in the theory literature of transcription, but also as an interpretive null model for huge classes of experimental data. To give a flavor for the use of these models, we provide several key case studies followed by a smorgasbord of citations which the reader is urged to consult. Aside from providing references, we decided to forego our own extensive efforts at using and scrutinizing thermodynamic models because we wanted to highlight the ubiquitous nature of such thinking beyond our own work. The original theory work of Ackers and Shea [10, 11] focused primarily on the example of phage lambda, itself already introduced non-mathematically in thermodynamic model format by Ptashne and collaborators [12]. Perhaps no example is more famous than the bacterial example of the *lac* operon and its synthetic variants. Müller-Hill and Oehler and coworkers made an impressive series of quantitative and rigorous studies of synthetic variants of the *lac* operon which in modern parlance we would view as having "tuned the knobs" of transcription such as the strength of DNA binding sites for Lac repressor, the copy number of the Lac repressor and even the length of the DNA loop formed by binding two sites simultaneously, done with exquisite single-base pair precision [19, 20]. In a large number of papers, Vilar and Leibler [13] and subsequently, Saiz and Vilar [21, 22, 23] have provided a corresponding theoretical study of this data (and much more). Kuhlman et al. used the tools of molecular biology to construct strains of *E. coli* such that they could explicitly test thermodynamic models [24](which they expertly modeled using thermodynamic models following their own earlier theory work showing how thermodynamic models could be used to dissect logic gates [14]). Similar studies in the context of the *ara* operon by Schleif and co-workers provided a picture of how DNA looping can be treated quantitatively within the confines of thermodynamic models [25, 26, 27]. We are strong advocates for those cases in which ultimately models of transcription are confronted with well-designed experiments that tune the same knobs that were controlled in the theoretical models. Another example of this kind of regulatory dissection for bacterial promoters is offered by work on MarA which activates transcription [28, 29]. One reason for skepticism concerning these apparent successes is the possibility that in some cases non-equilibrium and equilibrium models will "agree" on some particular set of data. To distinguish them may involve tuning some knob that has not yet been tuned. This paragraph only scratches the surface of the vast array of work based upon these models. A more detailed sense of the reach of this work can be gleaned by looking at the hundreds of citations of papers such as those of Buchler, Gerland and Hwa [14], Bintu et al. [30, 31], and Sherman and Cohen [15].

One of the most important measures of the "success" of thermodynamics is in their power to unify apparently quite distinct data in the form of data collapse. Two extremely impressive examples of such data collapse were explored in the context of chemotaxis (see Fig 5 of Keymer et al. [32]) and quorum sensing (see Fig 6 of Swem et al. [33]), where in both cases the activity of a signaling pathway was modeled using the equilibrium MWC model of allosteric receptors. The key finding is that the receptor activity for an entire suite of mutants could be collapsed onto one single master curve in the same way that for simple ligand-receptor binding (Hill function with Hill coefficient $n = 1$), if we plot $p_{\text{bound}}$ vs $c/K_d$ rather than $c$, we find that all ligand-receptor curves fall on one universal curve. For the chemotaxis and quorum sensing examples, the data collapse is much more subtle. Though we cannot consider this a bulletproof demonstration that the thermodynamic models are "right," they certainly provide a powerful, unifying and parameter-free predictive framework for thinking about experiments. In the context of transcription, similar data collapse was achieved featuring a very demanding parameter-free collapse of a broad array of experimental data in which binding site strength, transcription factor copy number and gene copy number were systematically varied (see Fig 4 of Weinert et al. [34]) and for these same constructs as a function of inducer concentration (see Fig 7(B) of Razo et al. [1]).

Before briefly turning to a survey of some of the shortcomings of the thermodynamic models, we present examples in which differential equations are used to model the dynamics of either mRNA or protein production and that implicitly feature thermodynamic models to describe gene regulation. Our key point here is to note that often these equations take the form (for example, Eq 5 in Cherry and Adler [35])

$$\frac{\mathrm{d}A}{\mathrm{d}t} = -\gamma A + f_{\text{production}}(A), \tag{S6}$$

where almost always the production term can be written in the form

$$f_{\text{production}}(A) = \frac{P(A)}{Q(A)}, \tag{S7}$$

where $P(A)$ and $Q(A)$ are polynomials. Further, in most instances, these rational functions are either of the phenomenological (or quasi-equilibrium) Hill form or appeal directly to Boltzmann states and weights. To be concrete, in two of the classic examples of synthetic biology, the genetic switch and the repressilator, the dynamical models were of the form described above [36, 37]. For example, for the genetic switch the production of the two species of repressor are written in dimensionless form (see Eq 1a and Eq 1b of Gardner et al. [36]) as

$$\begin{aligned} \frac{\mathrm{d}r_1}{\mathrm{d}\tau} &= -r_1 + \frac{\alpha}{1 + r_2^n}, \\ \frac{\mathrm{d}r_2}{\mathrm{d}\tau} &= -r_2 + \frac{\alpha}{1 + r_1^n}. \end{aligned} \tag{S8}$$

Here $r_1$ and $r_2$ are the dimensionless concentrations of the two mutually repressing repressors, time is measured in units of $\tau = \gamma t$ where $\gamma$ is the degradation rate and $\alpha$ is a dimensionless protein production rate. Our main point here is to note that although the words "thermodynamic model" or "equilibrium" are never used, the right-hand side of these equations is explicitly computing the probability of binding site occupancy by repressors.

We are hopeful that the subject of transcription will be held to the highest quantitative standards. We are excited by the many examples highlighted here (which only scratches the surface) and hope for a time when we have a deep and predictive understanding of all the genes in key model organisms and that these insights will serve as the basis for the study of non-model organisms such as redwood trees and blue whales. One route to such predictive understanding is the critical scrutiny offered by a dialogue between theory and experiment. In this paragraph, we note in passing a number of examples where it appears that the thermodynamic null models do not pass muster. One subject of intense effort over the last few decades is the study of DNA packing in eukaryotes and its implications for gene expression. As usual, the literature of this topic is immense. Intense debates have unfolded on the position of nucleosomes on genomic DNA with the conclusion likely that equilibrium models by themselves will *not* explain all the extant data [38, 39, 40]. The connection between expression and nucleosomal occupancy has been taken farther recently using single-cell methods with the result that the thermodynamic null model must be superseded by a more detailed model [41]. Similarly, in the context of the *Pho5* promoter in yeast, a quite amazing set of experiments was done to measure the occupancies of nucleosomes on this promoter. This data was analyzed using tens of thousands of models and the only models consistent with all of the data appear to require broken detailed balance [42, 43]. In a beautiful use of the MWC model highlighted above, Mirny worked out the probability that nucleosomes will be present essentially regulating some gene of interest and even went so far as to make an analogy with the Bohr effect in hemoglobin in which the post-translational modification of nucleosomes could be thought of as a kind of Bohr effect [44]. Though it took nearly a decade, it appears that this model is not sufficient to explain chromatin accessibility and gene expression [45]. We note that this example is common: to really make the comparison between theory and experiment often means that the data that exists is just not quite right to make the acid test (see the example of hemoglobin where, in our view, misuses of the MWC model led to the conclusion that the "model doesn't work" whereas in reality, it was rather that the set of states needed to be expanded to include other effectors [46, 47, 45]). This is also carefully explained in Chapter 7 of Ref. [48]. However, failure of the thermodynamic framework reaches well beyond the example of chromatin where it is clear that energy-consuming processes such as nucleosome remodeling can break detailed balance. Beautiful single-molecule experiments in *E. coli* revealed that even in the process of transcription factor binding to its DNA target, the results were not consistent with the thermodynamic framework [49]. Similarly, application of the thermodynamic modeling framework in the context of the *Pseudomonas aeruginosa* genes associated with the transcription factor BqsR have thus far been unsuccessful [50]. One particularly interesting example of the shortcomings of the thermodynamic approach was in the context of the glucocorticoid receptor where it was found that the rank ordering of gene expression did not scale with the rank ordering of binding site strength [51]. As with our list of "successes" this list of failures of the thermodynamic framework is superficial and incomplete. Further, we argue that to actually make claims of "right" or "wrong" requires diligent and often frustrating dialogue between theory and experiment where dedicated efforts need to be made to make sure that the same knobs are being tuned in both the theory and the corresponding experiments.

In summary, we would put the place of thermodynamic models in thinking about in vivo biochemistry (including the case of transcription) on par with other seemingly naive but incredibly productive null models such as the apparently crazy idea of a noninteracting electron gas to describe metals or the Ising model as a way to describe magnetic phenomena. In the context of transcription, though it is routine to critique thermodynamic models, we would argue that in fact, the thermodynamic models are just as plausible as the equally popular "two-state promoter" used so often to explain noise in transcription. Several beautiful examples of the use of the two-state framework are given in So at al. and Zenklusen et al. [52, 53]. For the purposes of the present paper, the thermodynamic model framework is a convenient null model which allows us to self-consistently generate synthetic datasets that can then be analyzed by the tools used to study real data with the added benefit that we "know the answer" from the outset. This appendix is a very superficial rendering of a vast subject. We hope at the least that it provides an entry into the literature which has used the so-called thermodynamic models and that attempts a balanced view of their successes and shortcomings. In our view, ultimately, the status of all models of transcription will only be really clarified by a painstaking dialogue between theory and experiment.

#### S2 Appendix Predicting the probability of RNA polymerase being bound using thermodynamic models

To estimate the expression levels of a gene with a given promoter, one key step is to calculate the probability of RNA polymerase (RNAP) being bound, which is needed in computing the rate of mRNA production using Eq 3. This can be done by building the so-called thermodynamic models of transcriptional regulation. In this appendix, we outline the protocol for building thermodynamic models, using a promoter that is constitutively expressed and a promoter with the simple repression architecture as examples.

On a broader level, the first step to build a thermodynamic model involves abstracting the genome into discrete microstates. As shown in Fig S1(A), our approach entails disregarding the three-dimensional topology of the genome and conceptualizing it as a linear sequence of discrete binding sites. RNAPs and transcription factors can bind to these sites in a number of configurations, each of which can be defined as a microstate. With this way of defining the microstates, we can then write down a general protocol for building thermodynamic models, which is shown in Fig S1(B). First, we need to identify all the relative promoter states. Second, we compute the energies for each of the states. Next, we need to calculate the multiplicity for each of the states. This is needed because each of the states encompass a range of possible configurations, and we need to count the total number of configurations associated with each state. Finally, we can compute the statistical weight of each state, which is the product of multiplicity and the Boltzmann weight calculated using the Boltzmann law of statistical mechanics. The statistical weight is in the form of

$$\omega_i = W\, e^{-\beta \varepsilon_i}, \tag{S9}$$

where $W$ is the multiplicity, $\beta = 1/k_B T$ with $k_B$ representing the Boltzmann constant and $T$ representing temperature, and $\varepsilon_i$ is the energy of the $i$-th state. The probability of any given state is then calculated by dividing $\omega_i$ by the partition function, which is the sum of the weights of all states.

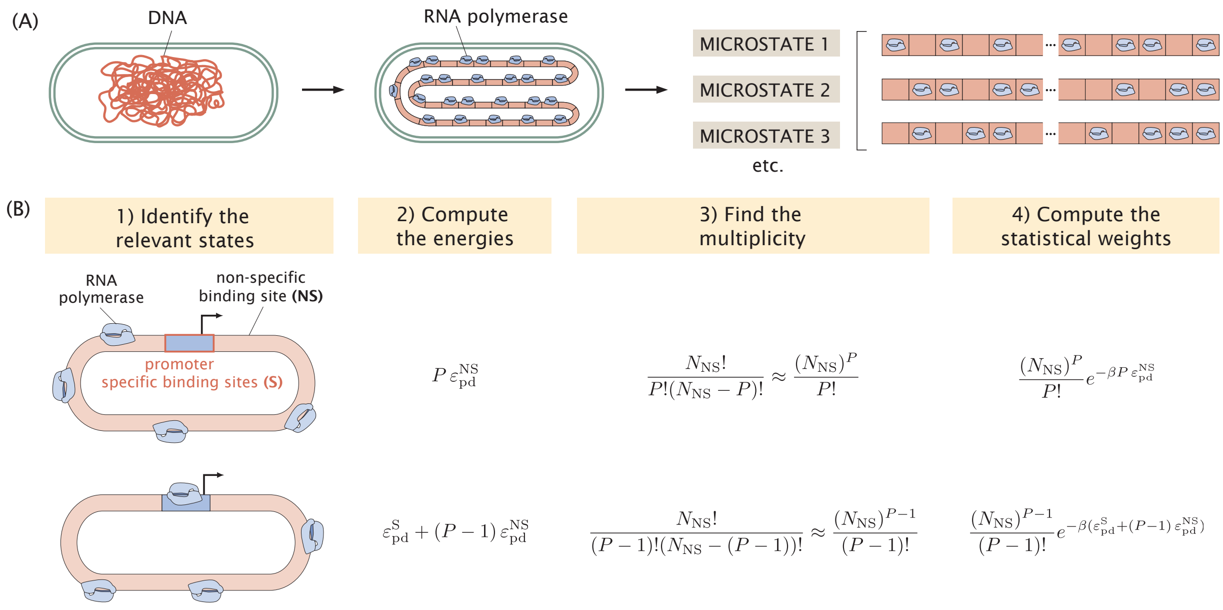


**Fig S1. Writing down thermodynamic models for transcriptional regulation.**
(A) Conceptualizing binding events along the genome as microstates. Each of the red boxes is a site along the genome to which RNAP and transcription factors can bind. Each of the possible configurations of binding is considered a microstate. (B) Protocol for writing down a thermodynamic model. The protocol involves four important steps. Firstly, all the relevant promoter states need to be identified. Secondly, the energies of each state is computed. Next, we compute the multiplicity, which tells us the number of possible configurations associated with each state. Finally, statistical weights are written down based on the energies and the multiplicity terms. For the energy terms $\varepsilon$, the superscript NS refers to non-specific binding, the superscript S refers to specific binding, the subscript pd refers to RNAP (p) binding to DNA (d).

To demonstrate how this protocol is used, let us consider a constitutive promoter, which is not regulated by any transcription factor. As shown in Fig S1(B), there are two possible states of binding: the state where one RNAP is bound to the promoter and the state where the promoter is empty. In the latter state, all RNAPs are bound to a non-specific binding site along the rest of the genome. Let us suppose that the specific binding energy at the promoter is $\varepsilon_{\mathrm{pd}}^{\mathrm{S}}$ and the non-specific binding energy along the rest of the genome is $\varepsilon_{\mathrm{pd}}^{\mathrm{NS}}$. If there are $P$ molecules of RNAPs in the system, then the energy of the empty promoter is $P\varepsilon_{\mathrm{pd}}^{\mathrm{NS}}$ and the energy when one RNAP is bound to the promoter and the remaining $(P-1)$ RNAPs are bound non-specifically is $\varepsilon_{\mathrm{pd}}^{\mathrm{S}} + (P-1)\varepsilon_{\mathrm{pd}}^{\mathrm{NS}}$.

Following the protocol, we next find the multiplicity of each promoter state. Let $N_{\mathrm{NS}}$ be the number of non-specific binding sites, then the multiplicity of the empty promoter state is given by

$$W_{\mathrm{NS}}(P, N_{\mathrm{NS}}) = \frac{N_{\mathrm{NS}}!}{P!(N_{\mathrm{NS}} - P)!} \approx \frac{(N_{\mathrm{NS}})^P}{P!}. \tag{S10}$$

The approximation in the second step holds true because $N_{\mathrm{NS}}$ is typically taken to be the length of the genome, which is on the order of $10^6$ for *E. coli*. Therefore, we have that $N_{\mathrm{NS}} \gg P$ and $\frac{N_{\mathrm{NS}}!}{(N_{\mathrm{NS}}-P)!} \approx (N_{\mathrm{NS}})^P$. We can use the same procedure to count the number of configurations associated with the state where RNAP is bound to the promoter. Since one RNAP molecule is bound to the promoter, there remain $(P-1)$ RNAP molecules that can bind to the non-specific binding sites in the rest of the genome. Therefore, the multiplicity of the RNAP-bound state is given by

$$W_{\mathrm{S}}(P-1, N_{\mathrm{NS}}) = \frac{N_{\mathrm{NS}}!}{(P-1)!(N_{\mathrm{NS}} - (P-1))!} \approx \frac{(N_{\mathrm{NS}})^{P-1}}{(P-1)!}. \tag{S11}$$

Having written down the energies and multiplicity terms of the two promoter states, we can compute the statistical weights of the states, which are given in the fourth column on Fig S1(B). Now we're ready to write down the probability of RNAP being bound for a constitutive promoter

$$p_{\mathrm{bound}} = \frac{\frac{(N_{\mathrm{NS}})^{P-1}}{(P-1)!} e^{-\beta\varepsilon_{\mathrm{pd}}^{\mathrm{S}}} e^{-\beta(P-1)\varepsilon_{\mathrm{pd}}^{\mathrm{NS}}}}{\frac{(N_{\mathrm{NS}})^{P}}{P!} e^{-\beta P\varepsilon_{\mathrm{pd}}^{\mathrm{NS}}} + \frac{(N_{\mathrm{NS}})^{P-1}}{(P-1)!} e^{-\beta\varepsilon_{\mathrm{pd}}^{\mathrm{S}}} e^{-\beta(P-1)\varepsilon_{\mathrm{pd}}^{\mathrm{NS}}}}, \tag{S12}$$

where $\varepsilon_{\mathrm{pd}}^{\mathrm{S}}$ is the binding energy of RNAP at the promoter and $\varepsilon_{\mathrm{pd}}^{\mathrm{NS}}$ is the binding energy of RNAP at the non-specific binding sites. In particular, we assume that the binding energy is same across all non-specific binding sites. To simplify the expression, we can multiply all the terms in the numerator and the denominator by $\frac{P!}{(N_{\mathrm{NS}})^P} e^{\beta P \varepsilon_{\mathrm{pd}}^{\mathrm{NS}}}$. This gives us

$$p_{\mathrm{bound}} = \frac{\frac{P}{N_{\mathrm{NS}}} e^{-\beta\Delta\varepsilon_{\mathrm{pd}}}}{1 + \frac{P}{N_{\mathrm{NS}}} e^{-\beta\Delta\varepsilon_{\mathrm{pd}}}}, \tag{S13}$$

where $\Delta\varepsilon_{\mathrm{pd}} = \varepsilon_{\mathrm{pd}}^{\mathrm{S}} - \varepsilon_{\mathrm{pd}}^{\mathrm{NS}}$ is the binding energy of the RNAP at the promoter relative to the binding energy at the non-specific binding sites.

This protocol can be easily extended to cases where the promoter is regulated by transcription factors. The simplest architecture that involves a transcription factor is where the promoter is regulated by a single repressor. As shown in Fig S2(B), for a promoter with the simple repression regulatory architecture, there are three possible state of binding: the state with an empty promoter, the state where RNAP is bound to the promoter, and the state where the repressor is bound to the promoter. Following the protocol above, we can write down the following statistical weights for each of the three states

$$\begin{aligned}
Z_{\text{empty promoter}} &= \frac{N_{\mathrm{NS}}!}{P!R!(N_{\mathrm{NS}} - P - R)!} e^{-\beta P\varepsilon_{\mathrm{pd}}^{\mathrm{NS}}} e^{-\beta R\varepsilon_{\mathrm{rd}}^{\mathrm{NS}}} && \text{(S14)}\\
&\approx \frac{(N_{\mathrm{NS}})^P}{P!} \frac{(N_{\mathrm{NS}})^R}{R!} e^{-\beta P\varepsilon_{\mathrm{pd}}^{\mathrm{NS}}} e^{-\beta R\varepsilon_{\mathrm{rd}}^{\mathrm{NS}}} && \text{(S15)}\\
Z_{\text{RNAP on promoter}} &= \frac{N_{\mathrm{NS}}!}{(P-1)!R!(N_{\mathrm{NS}} - (P-1) - R)!} e^{-\beta(P-1)\varepsilon_{\mathrm{pd}}^{\mathrm{NS}}} e^{-\beta R\varepsilon_{\mathrm{rd}}^{\mathrm{NS}}} e^{-\beta\varepsilon_{\mathrm{pd}}^{\mathrm{S}}} && \text{(S16)}\\
&\approx \frac{(N_{\mathrm{NS}})^{P-1}}{(P-1)!} \frac{(N_{\mathrm{NS}})^R}{R!} e^{-\beta(P-1)\varepsilon_{\mathrm{pd}}^{\mathrm{NS}}} e^{-\beta R\varepsilon_{\mathrm{rd}}^{\mathrm{NS}}} e^{-\beta\varepsilon_{\mathrm{pd}}^{\mathrm{S}}} && \text{(S17)}\\
Z_{\text{repressor on promoter}} &= \frac{N_{\mathrm{NS}}!}{P!(R-1)!(N_{\mathrm{NS}} - P - (R-1))!} e^{-\beta P\varepsilon_{\mathrm{pd}}^{\mathrm{NS}}} e^{-\beta(R-1)\varepsilon_{\mathrm{rd}}^{\mathrm{NS}}} e^{-\beta\varepsilon_{\mathrm{rd}}^{\mathrm{S}}} && \text{(S18)}\\
&\approx \frac{(N_{\mathrm{NS}})^P}{P!} \frac{(N_{\mathrm{NS}})^{R-1}}{(R-1)!} e^{-\beta P\varepsilon_{\mathrm{pd}}^{\mathrm{NS}}} e^{-\beta(R-1)\varepsilon_{\mathrm{rd}}^{\mathrm{NS}}} e^{-\beta\varepsilon_{\mathrm{rd}}^{\mathrm{S}}} && \text{(S19)}
\end{aligned}$$

where $N_{\mathrm{NS}}$ is the number of non-specific binding sites; $P$ is the number of RNAP; $R$ is the number of repressors; $\Delta\varepsilon_{\mathrm{pd}}$ is the binding energy of the RNAP; $\varepsilon_{\mathrm{pd}}^{\mathrm{S}}$ and $\varepsilon_{\mathrm{pd}}^{\mathrm{NS}}$ are the specific binding energy of the

RNAP at the promoter and the binding energy of the RNAP at the non-specific binding site; $\varepsilon_{\mathrm{rd}}^{\mathrm{S}}$ and $\varepsilon_{\mathrm{rd}}^{\mathrm{NS}}$ are the specific binding energy of the repressor at the promoter and the binding energy of the repressor at the non-specific binding site. This allows us to write down the probability of RNAP binding as

$$p_{\text{bound}} = \frac{Z_{\text{RNAP on promoter}}}{Z_{\text{empty promoter}} + Z_{\text{RNAP on promoter}} + Z_{\text{repressor on promoter}}}. \tag{S20}$$

Again, we simplify the expression by multiplying both the numerator and the denominator by $\frac{P!}{(N_{\mathrm{NS}})^P}\frac{R!}{(N_{\mathrm{NS}})^R}e^{\beta\varepsilon_{\mathrm{pd}}^{\mathrm{NS}}}e^{\beta\varepsilon_{\mathrm{rd}}^{\mathrm{NS}}}$. This gives us the following expression for the probability of RNAP being bound

$$p_{\text{bound}} = \frac{\frac{P}{N_{\mathrm{NS}}}e^{-\beta\Delta\varepsilon_{\mathrm{pd}}}}{1 + \frac{P}{N_{\mathrm{NS}}}e^{-\beta\Delta\varepsilon_{\mathrm{pd}}} + \frac{R}{N_{\mathrm{NS}}}e^{-\beta\Delta\varepsilon_{\mathrm{rd}}}}, \tag{S21}$$

where $\Delta\varepsilon_{\mathrm{pd}} = \varepsilon_{\mathrm{pd}}^{\mathrm{S}} - \varepsilon_{\mathrm{pd}}^{\mathrm{NS}}$ and $\Delta\varepsilon_{\mathrm{rd}} = \varepsilon_{\mathrm{rd}}^{\mathrm{S}} - \varepsilon_{\mathrm{rd}}^{\mathrm{NS}}$ are the binding energies of the RNAP and the repressors at the promoter relative to their non-specific binding energies. Here, the weak promoter approximation is often made, which states that the RNAP binding state has a much lower Boltzmann weight compared to the repressor binding site. Therefore, the expression can often be simplified to

$$p_{\text{bound}} = \frac{\frac{P}{N_{\mathrm{NS}}}e^{-\beta\Delta\varepsilon_{\mathrm{pd}}}}{1 + \frac{R}{N_{\mathrm{NS}}}e^{-\beta\Delta\varepsilon_{\mathrm{rd}}}}. \tag{S22}$$

It is important to note that when the expressions for $p_{\text{bound}}$ are used to predict the expression levels of promoter variants in an MPRA library, the energy terms $\varepsilon_i$ are calculated for each promoter variant by mapping binding site sequences to energy matrices. The procedure for calculating the total energies is explained in Sec 1.1 and illustrated in Fig 2(A).

### S3 Appendix States-and-weights models for common regulatory architectures

There are six common regulatory architectures for promoters in *E. coli*. In S2 Appendix, we have written down $p_{\text{bound}}$, the probability that the RNAP is bound to the promoter, for a constitutively expressed promoter and a promoter with the simple repression regulatory architecture. Based on the states-and-weights diagrams shown in Fig S2 and using the same protocol introduced in S2 Appendix, we can write down $p_{\text{bound}}$ the remaining four common regulatory architectures [30].

For a promoter with the simple activation regulatory architecture, the states-and-weights diagram is shown in Fig S2(C), and the probability of RNAP being bound is given by

$$p_{\text{bound}} = \frac{\frac{P}{N_{\mathrm{NS}}}e^{-\beta\Delta\varepsilon_{\mathrm{pd}}} + \frac{P}{N_{\mathrm{NS}}}\frac{A}{N_{\mathrm{NS}}}e^{-\beta(\Delta\varepsilon_{\mathrm{pd}}+\Delta\varepsilon_{\mathrm{ad}})}\omega_{\mathrm{ap}}}{1 + \frac{P}{N_{\mathrm{NS}}}e^{-\beta\Delta\varepsilon_{\mathrm{pd}}} + \frac{A}{N_{\mathrm{NS}}}e^{-\beta\Delta\varepsilon_{\mathrm{rd}}} + \frac{P}{N_{\mathrm{NS}}}\frac{A}{N_{\mathrm{NS}}}e^{-\beta(\Delta\varepsilon_{\mathrm{pd}}+\Delta\varepsilon_{\mathrm{ad}})}\omega_{\mathrm{ap}}}, \tag{S23}$$

where $N_{\mathrm{NS}}$ is the number of non-specific binding sites; $P$ is the number of RNAP; $A$ is the number of activators; $\Delta\varepsilon_{\mathrm{pd}}$ is the binding energy of the RNAP; $\Delta\varepsilon_{\mathrm{ad}}$ is the binding energy of the activator; $\omega_{\mathrm{a_1a_2}}$ is the interaction energy between the activator and the RNAP.

For a promoter with the repression-activation regulatory architecture, the states-and-weights diagram is shown in Fig S2(D), and the probability of RNAP being bound is given by

$$p_{\text{bound}} = \frac{\frac{P}{N_{\mathrm{NS}}}e^{-\beta\Delta\varepsilon_{\mathrm{pd}}} + \frac{P}{N_{\mathrm{NS}}}\frac{A}{N_{\mathrm{NS}}}e^{-\beta(\Delta\varepsilon_{\mathrm{pd}}+\Delta\varepsilon_{\mathrm{ad}})}\omega_{\mathrm{ap}}}{1 + \frac{P}{N_{\mathrm{NS}}}e^{-\beta\Delta\varepsilon_{\mathrm{pd}}} + \frac{R}{N_{\mathrm{NS}}}e^{-\beta\Delta\varepsilon_{\mathrm{rd}}} + \frac{A}{N_{\mathrm{NS}}}e^{-\beta\Delta\varepsilon_{\mathrm{ad}}} + \frac{P}{N_{\mathrm{NS}}}\frac{A}{N_{\mathrm{NS}}}e^{-\beta(\Delta\varepsilon_{\mathrm{pd}}+\Delta\varepsilon_{\mathrm{ad}})}\omega_{\mathrm{ap}}}, \tag{S24}$$

where $N_{\mathrm{NS}}$ is the number of non-specific binding sites; $P$ is the number of RNAP; $R$ is the number of repressors; $A$ is the number of activators; $\Delta\varepsilon_{\mathrm{pd}}$ is the binding energy of the RNAP; $\Delta\varepsilon_{\mathrm{rd}}$ is the binding energy of the repressor; $\Delta\varepsilon_{\mathrm{ad}}$ is the binding energy of the activator; $\omega_{\mathrm{a_1a_2}}$ is the interaction energy between the activator and the RNAP.

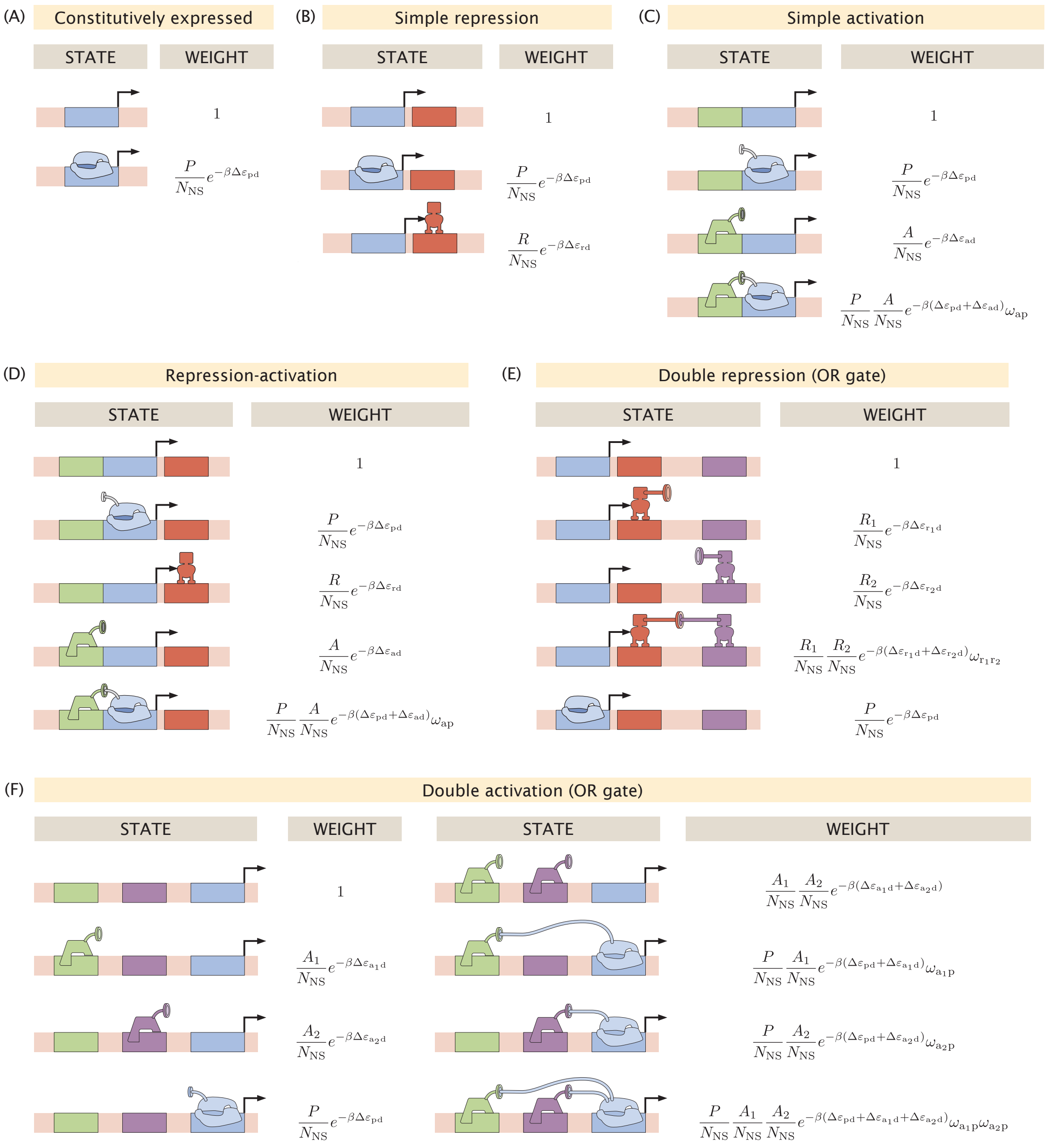


**Fig S2. States-and-weights models for common regulatory architectures.** In all the diagrams, $P$ represents the number of RNAP; $R$ represents the number of repressors; $A$ represents the number of activators; $N_{\mathrm{NS}}$ represents the number of non-specific binding sites; $\Delta\varepsilon_{\mathrm{pd}}$ represents the binding energy of the RNAP; $\Delta\varepsilon_{\mathrm{rd}}$ represents the binding energy of the repressor; $\Delta\varepsilon_{\mathrm{ad}}$ represents the binding energy of the activator; $\omega_{\mathrm{ab}} = e^{-\beta\varepsilon_{\mathrm{int}}}$ represents the interaction energy between a and b. (A) States-and-weights model for a promoter that is constitutively expressed. (B) States-and-weights model for a promoter under the simple repression regulatory architecture. (C) States-and-weights model for a promoter under the simple activation regulatory architecture. (D) States-and-weights model for a promoter under the repression-activation regulatory architecture. (E) States-and-weights model for a promoter under the double repression regulatory architecture with OR logic. (F) States-and-weights model for a promoter under the double activation regulatory architecture with OR logic.

Let $r_1 = \frac{R_1}{N_{\rm NS}} e^{-\beta \Delta\varepsilon_{r_1 d}}$, $r_2 = \frac{R_2}{N_{\rm NS}} e^{-\beta \Delta\varepsilon_{r_2 d}}$, and $p = \frac{P}{N_{\rm NS}} e^{-\beta \Delta\varepsilon_{pd}}$. Then, for a promoter with the double repression regulatory architecture under OR logic, the states-and-weights diagram is shown in Fig S2(E), and the probability of RNAP being bound is given by

$$p_{\rm bound} = \frac{p}{1 + r_1 + r_2 + r_1 r_2 \omega_{\rm r_1 r_2} + p}, \tag{S25}$$

where $N_{\rm NS}$ is the number of non-specific binding sites; $P$ is the number of RNAP; $R_1$ is the number of the first repressor; $R_2$ is the number of the second repressor; $\Delta\varepsilon_{\rm pd}$ is the binding energy of the RNAP; $\Delta\varepsilon_{\rm r_1 d}$ is the binding energy of the first repressor; $\Delta\varepsilon_{\rm r_2 d}$ is the binding energy of the second repressor; $\omega_{\rm r_1 r_2}$ is the interaction energy between the two repressors. The states-and-weights diagram of the AND-logic double repression regulatory architecture is shown in Fig 7(A). In this case, the probability of RNAP being bound is given by

$$p_{\rm bound} = \frac{p + r_1 p + r_2 p}{1 + r_1 + r_2 + r_1 r_2 \omega_{\rm r_1 r_2} + p + r_1 p + r_2 p}, \tag{S26}$$

Let $a_1 = \frac{A_1}{N_{\rm NS}} e^{-\beta \Delta\varepsilon_{a_1 d}}$, $a_2 = \frac{A_2}{N_{\rm NS}} e^{-\beta \Delta\varepsilon_{a_2 d}}$, and $p = \frac{P}{N_{\rm NS}} e^{-\beta \Delta\varepsilon_{pd}}$. Then, for a promoter with the double repression regulatory architecture under OR logic, the states-and-weights diagram is shown in Fig S2(F), and the probability of RNAP being bound is given by

$$p_{\rm bound} = \frac{p + a_1 p \omega_{a_1 p} + a_2 p \omega_{a_2 p} + a_1 a_2 p \omega_{a_1 p} \omega_{a_2 p}}{1 + a_1 + a_2 + p + a_1 a_2 \omega_{a_1 a_2} p + a_1 p \omega_{a_1 p} + a_2 p \omega_{a_2 p} + a_1 a_2 p \omega_{a_1 p} \omega_{a_2 p}}, \tag{S27}$$

where $N_{\rm NS}$ is the number of non-specific binding sites; $P$ is the number of RNAP; $A_1$ is the number of the first activator; $A_2$ is the number of the second activator; $\Delta\varepsilon_{\rm pd}$ is the binding energy of the RNAP; $\Delta\varepsilon_{\rm a_1 d}$ is the binding energy of the first activator; $\Delta\varepsilon_{\rm a_2 d}$ is the binding energy of the second activator; $\omega_{\rm a_1 p}$ is the interaction energy between the first activator and the RNAP; $\omega_{\rm a_1 a_2}$ is the interaction energy between the second activator and the RNAP. The states-and-weights diagram of the AND-logic double activation regulatory architecture is shown in Fig S8(A). In this case, the probability of RNAP being bound is given by

$$p_{\rm bound} = \frac{p + a_1 p \omega_{a_1 p} + a_2 p \omega_{a_2 p} + a_1 a_2 p \omega_{a_1 p} \omega_{a_2 p} \omega_{a_1 a_2}}{1 + a_1 + a_2 + p + a_1 a_2 \omega_{a_1 a_2} p + a_1 p \omega_{a_1 p} + a_2 p \omega_{a_2 p} + a_1 a_2 p \omega_{a_1 p} \omega_{a_2 p} \omega_{a_1 a_2}}, \tag{S28}$$

where $\omega_{\rm a_1 a_2}$ is the interaction energy between the two activators.

#### S4 Appendix Definition of probability distributions in the calculation of mutual information

In order to build an information footprint from data, we need to calculate the mutual information between expression levels and the base identity at each position in the sequence, which is defined as

$$I_i = \sum_b \sum_\mu {\rm Pr}_i(b, \mu) \log_2 \left( \frac{{\rm Pr}_i(b, \mu)}{{\rm Pr}_i(b) \, {\rm Pr}(\mu)} \right), \tag{S29}$$

where $b$ represents base identity and $\mu$ represents expression levels.

As shown in Fig S3(A) and S3(C), we find that binding sites have a higher signal-to-noise ratio in information footprints when the coarse grained approach is taken

$$b = \begin{cases} 0, & \text{if the base is mutated,} \\ 1, & \text{if the base is wild type.} \end{cases} \tag{S30}$$

On the other hand, to obtain a distribution of expression levels, sequencing counts are binned into $N$ bins, where the bins are chosen such there is an equal number of sequences in each bin. Again, we observe the highest signal-to-noise ratio for $N = 2$, and see a continuous decrease with increasing number of bins. As

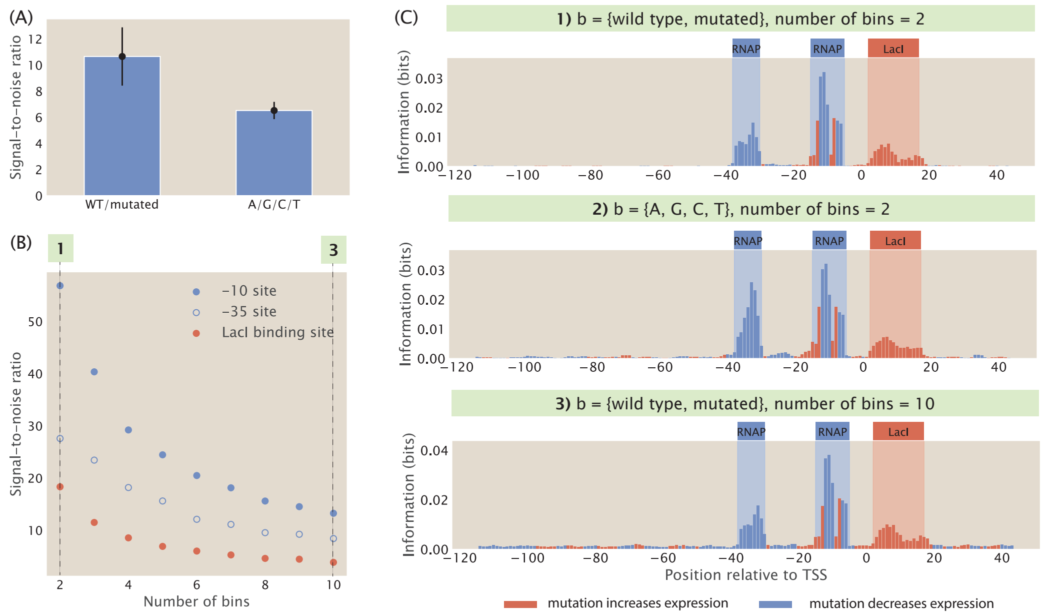


**Fig S3. Definition of probability distributions in the calculation of mutual information.** (A) Using a probability distribution of the four bases leads to a reduced signal-to-noise ratio in the information footprint. The heights of the bars are the average signal-to-noise ratios calculated from the information footprints of 20 synthetic datasets with the repression-activation architecture. (B) Signal-to-noise ratio decreases when the number of bins increases. Each data point is the mean of average mutual information across 20 synthetic datasets with the corresponding number of bins. The numbered labels correspond to footprints in (C). (C) Choosing different probability distributions to calculate the information footprint for a synthetic dataset with the repression-activation genetic architecture. The top footprint uses a probability distribution of wild-type and mutated bases and uses 2 bins to calculate the probability distribution for expression levels. The middle footprint uses a probability distribution of the four bases (A, G, C, T) and uses 2 bins to calculate the probability distribution for expression levels. The bottom footprint uses a probability distribution of wild-type and mutated bases and uses 10 bins to calculate the probability distribution for expression levels.

shown in Fig S3(B) and S3(C), we observe that signal-to-noise ratio is higher when fewer bins are used to partition expression levels.

These observations may be explained by the fact that increasing the number of states or bins increases the level of noise. When there are more states or more bins, fewer sequences will be present in each bin. This amplifies the hitch-hiking effects discussed in Sec 1.3, leading to a higher level of noise. In addition, since the boundaries between the bins are artificially set, a sequence may be randomly grouped into the N-th bin rather than the adjacent (N-1)-th and (N+1)-th bins simply because the boundaries are set a particular level. This randomness occurs in both the marginal probability distribution for $\mu$ and the joint probability distribution, resulting in noise that is increased when more bins are added.

### S5 Appendix Analytical calculation of information footprint

To better understand how mutations in the binding sites create signals in the information footprint, we derive the information footprint analytically for a constitutively expressed gene, i.e. the promoter of the gene only has a binding site for RNAP and is not bound by any transcription factors.

Consider a promoter region where the RNAP binding site is $l$ base pairs long and the probability of mutation at each site is $\theta$. Furthermore, the binding energy of RNAP to the wild-type sequence is denoted by $\Delta\varepsilon$ and we assume that at each position within the binding site, a mutation comes with cost $\Delta\Delta\varepsilon$ to the binding energy. Therefore, if there are $m$ mutations in the binding site, the total binding energy between

RNAP and the mutant binding site is $\Delta\varepsilon + m\Delta\Delta\varepsilon$.

In a sufficiently large data set, the ratio of sequences with mutation at position $i$ is given by

$$\Pr{}_i(b) = \begin{cases} 1-\theta, & \text{if } b = 0 \\ \theta, & \text{if } b = 1. \end{cases} \tag{S31}$$

Next, we determine $\Pr(\mu)$. As before, we define $\Pr(\mu)$ as the probability that a given sequence leads to high expression levels or low expression levels. To predict expression levels, we begin by calculating $p_{\text{bound}}$ for each promoter variant. Since the gene is constitutively expressed, the probability of RNAP binding is given by

$$p_{\text{bound}} = \frac{\frac{P}{N_{\text{NS}}} e^{-\beta(\Delta\varepsilon + m\Delta\Delta\varepsilon)}}{1 + \frac{P}{N_{\text{NS}}} e^{-\beta(\Delta\varepsilon + m\Delta\Delta\varepsilon)}}. \tag{S32}$$

As derived in Eq 5, the steady state copy number of mRNAs is proportional to the probability of the RNAP bound state. Therefore, expression level is only dependent on the number of mutations in the RNAP binding site.

The probability distribution for the number of mutations in the RNAP binding site can be expressed using the binomial distribution, where the probability of $k$ mutations in the binding site is given by

$$\Pr(m = k; l, \theta) = \binom{l}{k} \theta^k (1-\theta)^{l-k}. \tag{S33}$$

As illustrated in Fig S4, since expression levels are solely determined by the number of mutations in the binding site, and sequences are binned by expression levels to obtain $P(\mu)$, there is a threshold number of mutations, $m^*$, where sequences with $m^*$ or more than $m^*$ mutations fall into the lower expression bin. Hence, $P(\mu)$ is given by

$$\Pr(\mu) = \begin{cases} \Pr(m \geq m^*; l, \theta) = \sum\limits_{k=m^*}^{l} \binom{l}{k} \theta^k (1-\theta)^{l-k}, & \text{if } \mu = 0 \\ \Pr(m < m^*; l, \theta) = 1 - \Pr(m \geq m^*; l, \theta), & \text{if } \mu = 1. \end{cases} \tag{S34}$$

Finally, we determine the expression for $\Pr_i(b, \mu)$. To do this, we consider two cases, one where the position $i$ is outside of the RNAP binding site and one where the position $i$ is within the RNAP binding site. If $i$ is not in the RNAP binding site $\mathcal{B}$, then a mutation would have no effect on the expression levels, therefore

$$\Pr{}_{i \notin \mathcal{B}}(b, \mu) = \begin{cases} (1-\theta) \cdot \Pr(m \geq m^*; l, \theta), & \text{if } b = 0 \text{ and } \mu = 0 \\ (1-\theta) \cdot \Pr(m < m^*; l, \theta), & \text{if } b = 0 \text{ and } \mu = 1 \\ \theta \cdot \Pr(m \geq m^*; l, \theta), & \text{if } b = 1 \text{ and } \mu = 0 \\ \theta \cdot \Pr(m < m^*; l, \theta), & \text{if } b = 1 \text{ and } \mu = 1. \end{cases} \tag{S35}$$

Having derived all the marginal probability distributions and the joint probability distributions, we can then write down mutual information at a non-binding site and at a binding site. If position $i$ is outside the RNAP binding site, then

$$\begin{aligned} I_i = & (1-\theta) \cdot \Pr(m \geq m^*; l, \theta) \log_2 \frac{(1-\theta) \cdot \Pr(m \geq m^*; l, \theta)}{(1-\theta) \cdot \Pr(m \geq m^*; l, \theta)} \\ & + (1-\theta) \cdot \Pr(m < m^*; l, \theta) \log_2 \frac{(1-\theta) \cdot \Pr(m < m^*; l, \theta)}{(1-\theta) \cdot \Pr(m < m^*; l, \theta)} \\ & + \theta \cdot \Pr(m \geq m^*; l, \theta) \log_2 \frac{\theta \cdot \Pr(m \geq m^*; l, \theta)}{\theta \cdot \Pr(m \geq m^*; l, \theta)} \\ & + \theta \cdot \Pr(m < m^*; l, \theta) \log_2 \frac{\theta \cdot \Pr(m < m^*; l, \theta)}{\theta \cdot \Pr(m < m^*; l, \theta)}. \end{aligned} \tag{S36}$$

We can see that $I_i = 0$ since the fractions within the logarithms all cancel out to be 1. This is because the joint probability $\Pr_i(b, \mu)$ for bases outside the binding site is simply given by the product of the marginal distributions,

$$\Pr_{i \notin \mathcal{B}}(b, \mu) = \Pr_{i \notin \mathcal{B}}(\mu) \Pr_{i \notin \mathcal{B}}(b). \tag{S37}$$

If the position $i$ is in the RNAP binding site, the calculation for $\Pr_i(b, \mu)$ is more complex. As illustrated in Fig S4, if the position $i$ has wild-type base identity, then the sequence would have low expression levels if there are more than $m^*$ mutations in the remaining $l-1$ bases in the RNAP binding site and the sequence would have high expression levels if there are less than $m^*$ mutations in the remaining $l-1$ bases in the RNAP binding site. On the other hand, if the position $i$ is mutated, then the sequence would have low expression levels if there are more than $m^*-1$ mutations in the remaining $l-1$ bases and the sequence would have high expression levels if there are less than $m^*-1$ mutations in the remaining $l-1$ bases. Taken together, we can write down the joint probability distribution as

$$\Pr_{i \in \mathcal{B}}(b, \mu) = \begin{cases} (1-\theta) \cdot \Pr(m \geq m^*; l-1, \theta), & \text{if } b = 0 \text{ and } \mu = 0 \\ (1-\theta) \cdot \Pr(m < m^*; l-1, \theta), & \text{if } b = 0 \text{ and } \mu = 1 \\ \theta \cdot \Pr(m \geq m^*-1; l-1, \theta), & \text{if } b = 1 \text{ and } \mu = 0 \\ \theta \cdot \Pr(m < m^*-1; l-1, \theta), & \text{if } b = 1 \text{ and } \mu = 1. \end{cases} \tag{S38}$$

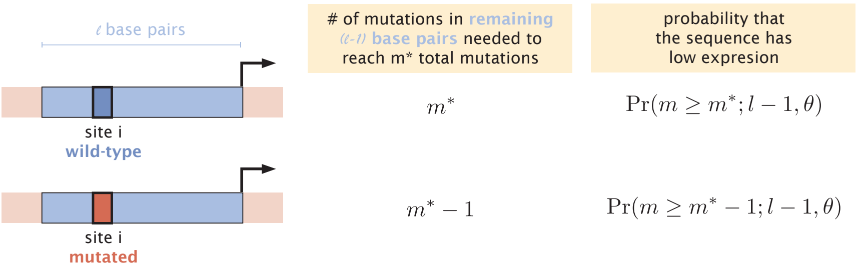


**Fig S4. Calculating number of mutations needed to reach the threshold between low expression and high expression bins.** The joint probability distribution at site $i$ is a product of the probability that site $i$ is mutated or wild type and the probability that the sequence has high or low expression level. Since the expression of the sequence depends on the presence of a mutation at site $i$, we need to consider two different cases in order to calculate the probability of expression. In the case where site $i$ has wild-type base identity, there need to be $m^*$ mutations outside of site $i$ in the RNAP binding site for the sequence to reach the threshold $m^*$. Therefore, the probability that the sequence has low expression is $\Pr(m \geq m^*; l-1, \theta)$. On the other hand, in the case where site $i$ is mutated, since one mutation is known to exist, there only need to be $m^*-1$ mutations outside of site $i$ in the RNAP binding site for the sequence to reach the threshold. In this case, the probability that the sequence has low expression is $\Pr(m \geq m^*-1; l-1, \theta)$.

In this case, the joint distribution does not factor into the marginal distributions,

$$\Pr_{i \in \mathcal{B}}(b, \mu) \neq \Pr_{i \in \mathcal{B}}(\mu) \Pr_{i \in \mathcal{B}}(b), \tag{S39}$$

and therefore, mutual information has to be larger than zero, $I_i > 0$, clearly distinguishing positions that are within the binding site from positions outside. Specifically,

$$\begin{aligned}
I_i =&(1-\theta)\cdot \Pr(m \geq m^*; l-1, \theta) \log_2 \frac{(1-\theta)\cdot \Pr(m \geq m^*; l-1,\theta)}{(1-\theta)\cdot \Pr(m \geq m^*; l,\theta)} \\
&+ (1-\theta)\cdot \Pr(m < m^*; l-1, \theta) \log_2 \frac{(1-\theta)\cdot \Pr(m < m^*; l-1,\theta)}{(1-\theta)\cdot \Pr(m < m^*; l,\theta)} \\
&+ \theta\cdot \Pr(m \geq m^*-1; l-1, \theta) \log_2 \frac{\theta\cdot \Pr(m \geq m^*-1; l-1,\theta)}{\theta\cdot \Pr(m \geq m^*; l,\theta)} \\
&+ \theta\cdot \Pr(m < m^*-1; l-1, \theta) \log_2 \frac{\theta\cdot \Pr(m < m^*-1; l-1,\theta)}{\theta\cdot \Pr(m < m^*; l,\theta)} \\
=&(1-\theta)\cdot \Pr(m \geq m^*; l-1, \theta) \log_2 \frac{\Pr(m \geq m^*; l-1,\theta)}{\Pr(m \geq m^*; l,\theta)} \\
&+ (1-\theta)\cdot \Pr(m < m^*; l-1, \theta) \log_2 \frac{\Pr(m < m^*; l-1,\theta)}{\Pr(m < m^*; l,\theta)} \\
&+ \theta\cdot \Pr(m \geq m^*-1; l-1, \theta) \log_2 \frac{\Pr(m \geq m^*-1; l-1,\theta)}{\Pr(m \geq m^*; l,\theta)} \\
&+ \theta\cdot \Pr(m < m^*-1; l-1, \theta) \log_2 \frac{\Pr(m < m^*-1; l-1,\theta)}{\Pr(m < m^*; l,\theta)}.
\end{aligned} \tag{S40}$$

### S6 Appendix Recovering binding site signal under extreme mutation rates

As we have shown in Sec 1.2, when the rate of mutation in the mutant library is low, we lose the signal at the RNAP binding site. We hypothesize that this is because RNAP binds weakly at the promoter. We generated a synthetic dataset that has low mutation rate but stronger binding energy at the RNAP binding site. As shown in Fig S5(A), the information footprint built from this dataset has a much higher level of mutual information at the RNAP binding site compared to the information footprint built from a dataset with the same mutation rate but weak RNAP binding energy, which supports our hypothesis.

We also showed that when the rate of mutation in the mutant library is high, there is low mutual information at the repressor binding site. Our hypothesis is that this is caused by the large effects of mutations on the repressor binding energy. We generated a synthetic dataset with high mutation rate while reducing the effect of mutation on binding energy by five fold. As shown in Fig S5(B), this allows us to recover the signal at the repressor binding site, which is also in line with our hypothesis.

### S7 Appendix Optimal mutation rate for various parameters

An essential part of designing libraries for MPRAs is the rate at which bases in the sequences are mutated. In Sort-Seq [54] and Reg-Seq [55], the mutation rate for promoter sequences is chosen as 0.1 per base. In Sec 1.2, we have calculated that this is on par with the optimal mutation rate for a promoter regulated by one transcription factor, given a specific set of parameters for copy numbers and binding energies of the transcription factor and RNAP. Here, we explore how the result for the optimal mutation rate depends on the specific choice of parameters.

In Eq 16, we have defined the optimal mutation rate as the rate where the Boltzmann weights of RNAP binding and transcription factor binding are equal. Here we explore how this mutation rate depends on the binding energy of RNAP $\Delta\varepsilon_{\text{pd}}$, the binding energy of the transcription factor $\Delta\varepsilon_{\text{rd}}$, the copy number of RNAP $P$, and the copy number of the transcription factor $R$.

For each set of parameters, we can solve Eq 16 numerically for the mutation rate that gives us $\kappa = 1$. In Fig S6, the optimal mutation rate is computed numerically when two of the parameters are varied while the others are kept constant. Increasing the binding energy or copy number of the transcription factor increases the mutation rate, while increasing the binding energy of RNAP and increasing the copy number of RNAP decreases the mutation rate. Changing the binding energy can have drastic effects on the mutation rate, in

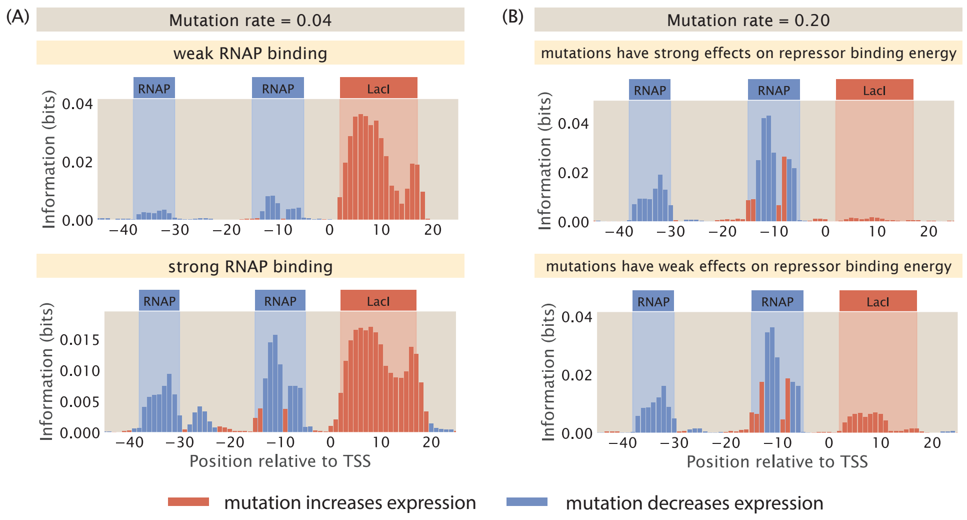


**Fig S5. Recovering signals from information footprints under extreme mutation rates.** (A) We generated two synthetic datasets with a mutation rate of 0.04 in the mutant library. In the first dataset, we set the RNAP binding energy $\Delta\varepsilon_{rd}$ to be $-5\ k_BT$, which is typical of RNAP binding at the wild type -10 and -35 binding sites. In the footprint produced from this dataset, there is low mutual information at the RNAP binding site due to the low mutation rate. On the other hand, in the second dataset, we increased $\Delta\varepsilon_{rd}$ to $-12\ k_BT$. This allows us to recover the signal at the RNAP binding site. (B) We generated two synthetic datasets with a mutation rate of 0.20. In the first dataset, we used the experimentally measured energy matrix for LacI at the O1 operator shown in Fig 2(B), where the average effect of mutations on binding energy is 2.24 $k_BT$. In the footprint produced from this dataset, there is low mutual information at the repressor binding site due to the high mutation rate. In the second dataset, we reduced the average effect of mutations five-fold and are able to recover the signal at the repressor binding site.

contrast to changes in copy numbers. This can be explained by the fact that the binding energies contributing exponentially to $\kappa$, while the copy numbers come into play as linear factors. For cases where the transcription factor bound state becomes very unlikely, e.g. in the cases of very weak binding of the transcription factor or very strong binding of the RNAP, there is no optimal mutation rate that can be found given the criteria in Eq 16. These regions can be found in Fig S6(B)-(D) as grey regions.

Binding sites and transcription factors come with widely different values for the parameters we have tested, e.g., the copy number of the activator CRP can be as high as about 500, while the copy number for an essential transcription factor DicA can be as low as 10 as measured in mass-spectrometry experiments [56]. The binding energy for the lac-repressor varies on the order of $6\,k_\mathrm{B}T$ ($-15.7\,k_\mathrm{B}T$ for the O1 operator and $-9.3\,k_\mathrm{B}T$ for the O3 operator [57]. Hence, it can be very beneficial to create a library of mutant sequences that contains sequences with different mutation rates in order to detect binding sites with these different parameters.

### S8 Appendix Changing transcription factor copy numbers for a double repression promoter under XOR gate

In Sec 2.2, we examined how changing transcription factor copy numbers affect the footprint for a double repression promoter under AND and OR logic gates. As discussed by Buchler et al. [14] and de Ronde et al. [58], only a limited number of logic gates are attainable through parameter variations, and a thermodynamic model can be written down for each of the possible logic gates. This means that our thermodynamic-model-based computational pipeline can be easily adapted to consider all possible types of interactions between transcription factors.

As an example, let us consider the exclusive-or (XOR) logic gate in a promoter regulated by two

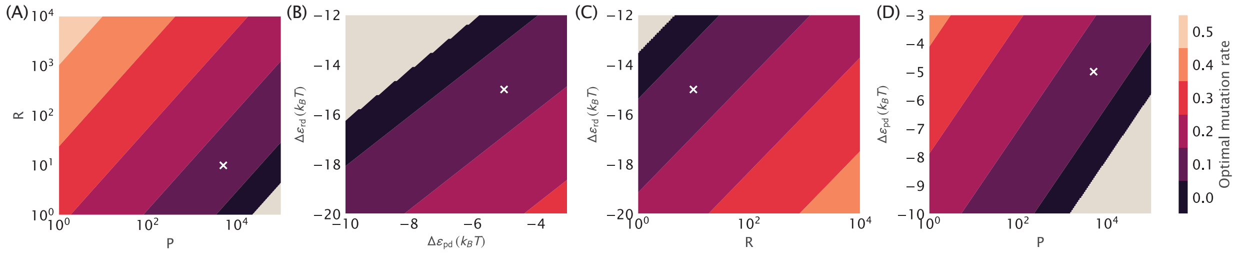


**Fig S6. Optimal mutation rate for single transcription factor binding.** Numerical solutions for the mutation rate calculated by finding the mutation rate that leads to equal Boltzmann weights between the RNAP bound state and the transcription factor bound state using Eq 16. The white crosses mark the standard set of parameters: $R = 10$, $P = 5000$, $\Delta\varepsilon_{\mathrm{rd}} = -15\,k_{\mathrm{B}}T$ and $\Delta\varepsilon_{\mathrm{pd}} = -15\,k_{\mathrm{B}}T$. Panels (A) to (D) vary 2 of the 4 parameters, while the other two stay constant. Gray color indicates regimes where no mutation rate can be found that fulfills the criteria of $\kappa = 1$.

repressors, which is another important and interesting logic gate other than the AND and OR gates. As illustrated in Fig S7(A), under the XOR gate, gene expression is repressed when only one of the repressors is present at high concentrations, but not when both of the repressors are present at high concentrations. One possible mechanism by which this may occur is if the interaction between the two repressors is repulsive. As shown in Fig S7(B) and S7(C), when the copy number of the second repressor is kept constant at 25 and the copy number of the first repressor is increased from 0 to 50, the signal at the first repression binding site increases and the signal at the second repressor binding site decreases. This behaviour is consistent with the definition of XOR logic gates and demonstrates that our computational pipeline can handle a diverse range of interaction regimes.

#### S9 Appendix Transcription factor knock-out under double activation

A double-activation promoter can also operate under an AND or an OR logic gate [14]. The states-and-weights diagram for a double-activation promoter is shown in Fig S8(A). Under AND logic, the two activators can interact both with the RNAP and with each other. This cooperativity leads to a further increase in expression levels. In contrast, under OR logic, the activators independently interact with RNAP and there is no cooperativity between them. We build synthetic datasets for an AND-logic and an OR-logic double-activation promoter. As shown in Fig S8(B) and S8(C), under AND logic, since cooperativity is at play, the signal at both $A_1$ and $A_2$ binding sites increases when $A_1$ is increased. On the other hand, under OR logic, the two activators act independently and there is competition between the signals at the two sites. When $A_1$ is increased, the signal at $A_1$ binding site correspondingly increases but the signal at $A_2$ binding site decreases.

#### S10 Appendix Changing inducer concentration for the inducible activator

In Sec 2.4, we discussed the effects of inducer concentration on the information footprints of a simple repression promoter with an inducible repressor. Similar effects can also be seen for a simple activation promoter with an inducible activator. One example of an inducible activator is CRP, which changes its conformation when bound to cyclic-AMP and thereby becomes more favorable to DNA binding [59]. Based on the states-and-weights diagram for such a promoter, which is shown in Fig S9(A), the probability of RNAP being bound is given by

$$p_{\mathrm{bound}} = \frac{p + p a_A \omega_A + p a_I \omega_I}{1 + p + a_A + a_I + p a_A \omega_A + p a_I \omega_I}, \tag{S41}$$

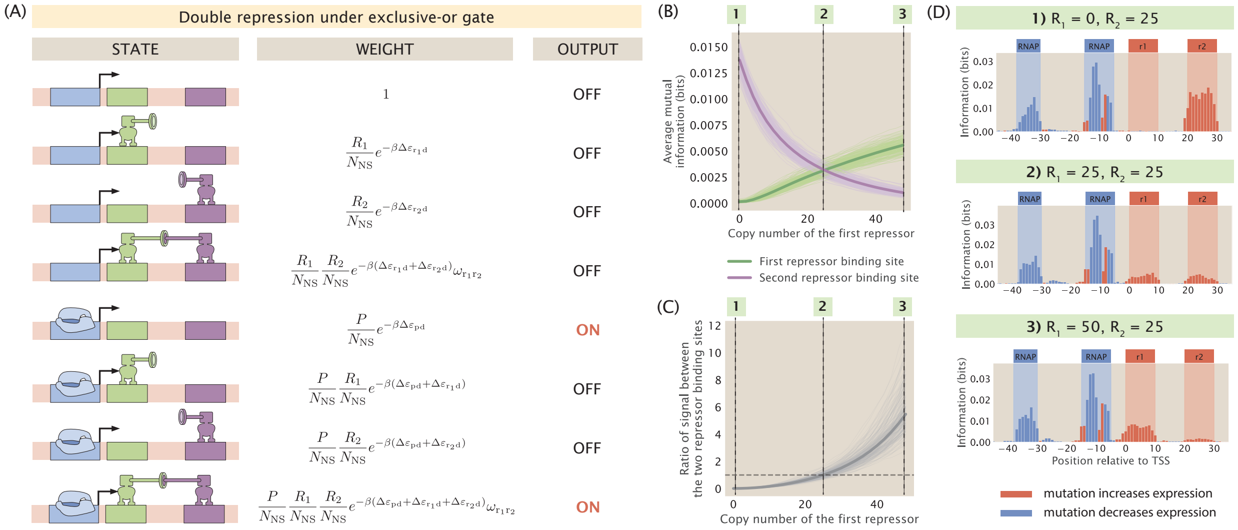


**Fig S7. Changing repressor copy number for a double-repression promoter under the XOR logic gate.** (A) States-and-weights diagram of a promoter with the double repression regulatory architecture and under the exclusive-or (XOR) gate. (B) Changes in the average mutual information at the two repressor binding sites when the copy number of the first repressor is increased. For the energy matrices of the repressors, the interaction energy between the repressor and a site is set to $0\,k_BT$ if the site has the wild type base identity and set to $1\,k_BT$ if the site has the mutant base identity. To enforce the XOR logic, the interaction energy between the repressors is set to $5\,k_BT$. 200 synthetic datasets are simulated and the trajectory for each of the synthetic dataset is shown as an individual light green or light purple curve. The average trajectories across all 200 synthetic datasets are shown as the bolded green curves and the bolded purple curves. The three numbered labels correspond to the information footprints shown in (D). (C) Ratio of average mutual information between the two repressor binding sites when the copy number of the first repressor changes. The individual trajectories (plotted in light grey) and mean trajectory (plotted in dark grey) are from the same 200 synthetic datasets used in generate the plot in (B). As expected, the ratio is equal to 1 when the copy number of the first repressor is equal to the copy number of the second repressor. (D) Representative information footprints with three different combination of repressor copy numbers.

where $p$ is the normalized weight of the RNAP bound state, $a_A$ is the normalized weight of the active activator bound state, and $a_I$ is the normalized weight of the inactive activator bound state. $\omega_A$ and $\omega_I$ account for the interaction energy between the RNAP and the active activator and the interaction energy between the RNAP and the inactive activator, respectively. The exact expressions for $p$, $a_A$, $a_I$, $\omega_A$ and $\omega_I$ are given in Fig S9(A).

To simplify the expression above, we determine the proportion of active and inactive activators with respect to the total number of activators. Similar to the case of simple repression, we calculate $p_{\text{active}}(c)$, which is the probability that the activator exists in the active conformation as a function of the inducer concentration, $c$. The different states of the activator can be modelled using the states-and-weights diagram shown in Fig S7(B). Here, we consider two types of cooperativity. The first type of cooperativity is between the two binding sites, where each ligand binding event changes the binding affinity of the unbound site. This is inherent to the classic MWC model and is already encoded in the terms $\omega_A$ and $\omega_I$ in Eq S41. The second type of cooperativity is between the two ligands, which accounts for the negative cooperativity of CRP in the inactive state. This is accounted for by the cooperative energy terms $\varepsilon_{\text{int}}^A$ and $\varepsilon_{\text{int}}^I$, which represent the interaction energies between the two ligands in the active and inactive states, respectively. Given the states-and-weights diagram, $p_{\text{active}}(c)$ is given by

$$p_{\text{active}}(c) = \frac{1 + \frac{c}{K_L^A} + \frac{c}{K_R^A} + \frac{c}{K_L^A}\frac{c}{K_R^A}e^{-\beta\varepsilon_{\text{int}}^A}}{1 + \frac{c}{K_L^A} + \frac{c}{K_R^A} + \frac{c}{K_L^A}\frac{c}{K_R^A}e^{-\beta\varepsilon_{\text{int}}^A} + e^{-2\beta\varepsilon_{AI}}(1 + \frac{c}{K_L^I} + \frac{c}{K_R^I} + \frac{c}{K_L^I}\frac{c}{K_R^I}e^{-\beta\varepsilon_{\text{int}}^I})}, \tag{S42}$$

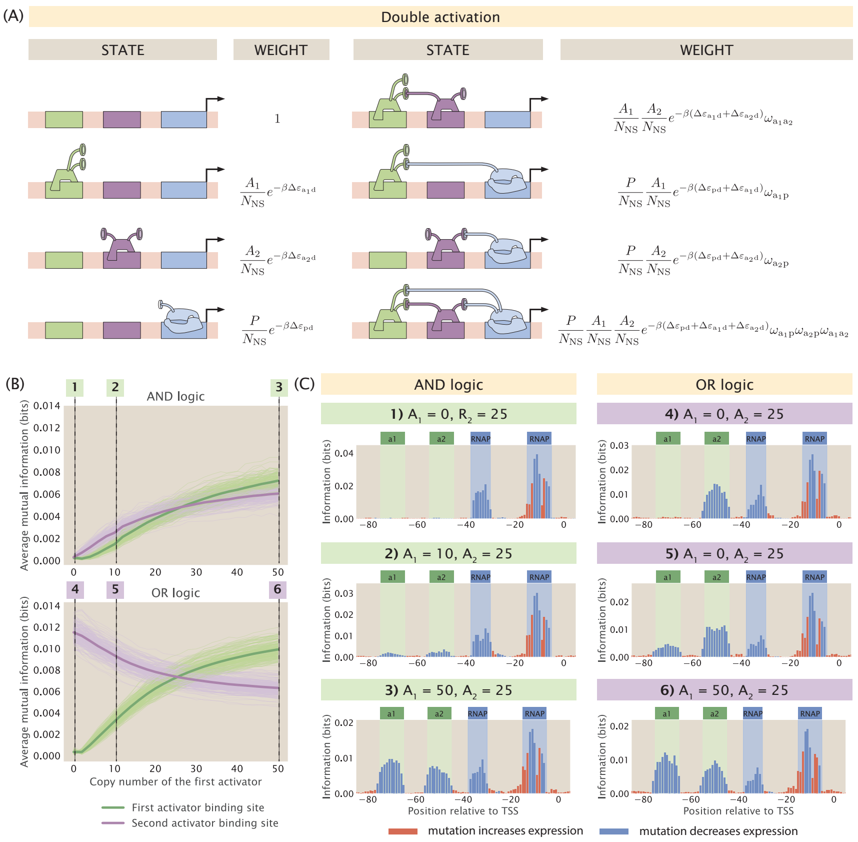


**Fig S8. Changing the copy number of activators in a double activation promoter.**
(A) States-and-weights diagram of a promoter with the double activation regulatory architecture. Under OR logic, the two activators do not exhibit coorperativity and $\omega_{\mathrm{a_1a_2}} = 0\ k_BT$. The states-and-weights diagram of a double activation promoter with OR logic is also shown in Fig S2(F). (B) Changing the copy number of the first activator under AND logic and OR logic affects the signal at both activator binding sites. The energy matrices of the activators are randomly generated in the same way as the energy matrices of the repressors in Fig 8. For the promoter with AND logic, the interaction energies between the activators and between the activator and the RNAP are set to -4 $k_BT$. For the promoter with OR logic, the interaction energies between the activators and between the activator and the RNAP are set to -7 $k_BT$. The higher interaction energy for the OR logic promoter is to ensure that there are similar levels of signal at the activator binding sites compared to the AND logic promoter. 200 synthetic datasets are simulated and the trajectory for each of the synthetic dataset is shown as an individual light green or light purple curve. The average trajectories across all 200 synthetic datasets are shown as the bolded green curves and the bolded purple curves.
(C) Representative information footprints of a double repression promoter under AND and OR logic.

where $K_L^A$ is the dissociation constant between the inducer and the left binding pocket of the active activator, $K_R^A$ is the dissociation constant between the inducer and the right binding pocket of the active activator, $K_L^I$ is the dissociation constant between the inducer and the left binding pocket of the inactive activator, and $K_R^I$ is the dissociation constant between the inducer and the right binding pocket of the inactive activator. With this expression, we can represent the number of active and inactive activators as $A_A = p_{\text{active}} A$ and $A_I = (1 - p_{\text{active}})A$. Therefore, we have that $a_A = p_{\text{active}} \frac{A}{N_{\text{NS}}} e^{-\beta \Delta \varepsilon_{ad}^A}$ and $a_I = (1 - p_{\text{inactive}}) \frac{A}{N_{\text{NS}}} e^{-\beta \Delta \varepsilon_{ad}^I}$.

We built synthetic datasets for a simple activation promoter with an inducible activator. As shown in Fig S9(C) and Fig S9(D), when the concentration of the inducer is increased, the average signal at the RNAP binding site increases, which corresponds to an increase in expression. At high inducer concentration, the activator is too strongly bound to be affected by mutations, and therefore the signal at the activator binding site is negligible.

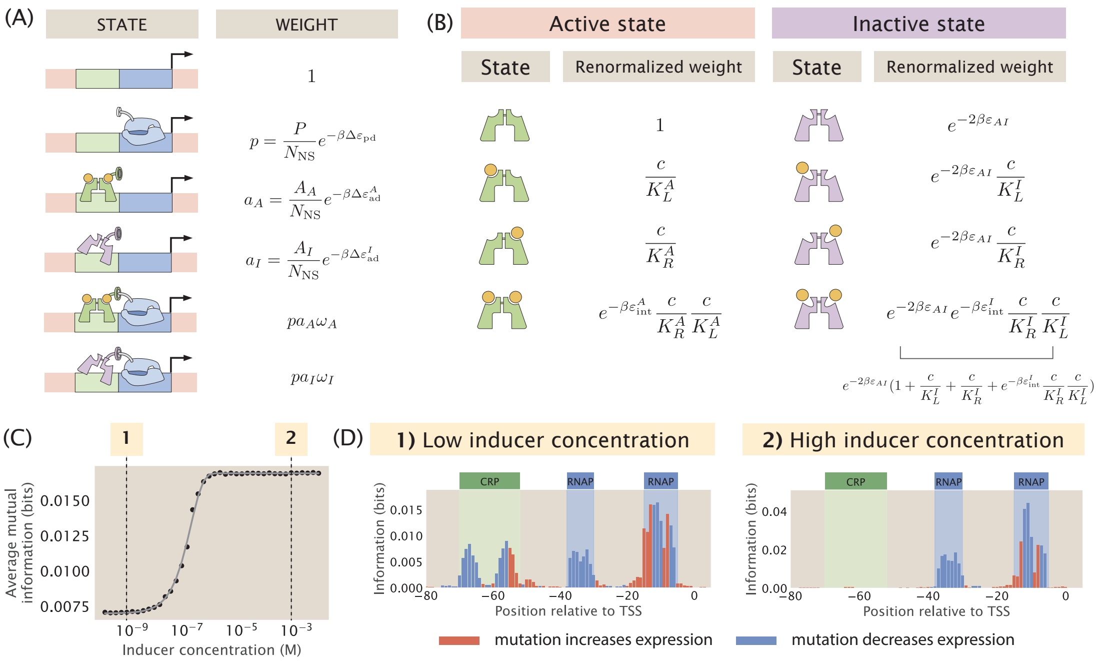


**Fig S9. Changing inducer concentration for the inducible activator.** (A) States-and-weights diagram for an inducible activator. In the diagram, $N_{\text{NS}}$ is the number of non-binding sites in the genome, $P$ is the copy number of the RNAP, $A_A$ is the copy number of active activators, $A_I$ is the copy number of inactive activators, $\Delta\varepsilon_{\text{pd}}$ is the binding energy of the RNAP, $\Delta\varepsilon^A_{\text{ad}}$ is the binding energy of the active activator, $\Delta\varepsilon^I_{\text{ad}}$ is the binding energy of the inactive activator. $\omega_A = e^{-\beta\varepsilon_{p,a_A}}$ and $\omega_I = e^{-\beta\varepsilon_{p,a_I}}$, where $\varepsilon_{p,a_A}$ is the interaction energy between the RNAP and the active activator and $\varepsilon_{p,a_I}$ is the interaction energy between the RNAP and the inactive activator. (B) States-and-weights diagram to calculate the probability that the activator is in the active state. (C) Average mutual information at the RNAP binding site increases as the inducer concentration increases. Here, we let $K^A_L = K^A_R = 3 \times 10^{-6}$ M, $K^I_L = K^A_R = 10^{-7}$ M, and $\Delta\varepsilon_{AI} = -3\ k_BT$ [59]. Each data point is the mean of average mutual information across 20 synthetic datasets with the corresponding inducer concentration. The numbered labels correspond to footprints in (D). (D) Representative information footprints with low inducer concentration ($10^{-9}$ M) and high inducer concentration ($10^{-3}$ M).

### S11 Appendix Noise from experimental procedures

In MPRAs such as Reg-Seq, the mutant library is grown up in culture. Once the cell culture is prepared, genomic DNA (gDNA) and mRNAs are extracted, the latter of which is used as a template in reverse transcription to make complementary DNA (cDNA). Afterwards, polymerase chain reation (PCR) is performed to amplify the reporter gene from the gDNA and cDNA. Finally, sequencing adapters are attached to the gDNA and cDNA. The gDNA and cDNA are then sequenced to obtain DNA and RNA counts for each sequence variant.

As illustrated in Fig S10(A), there are at least two possible sources of experimental noise. Firstly, PCR amplification is a stochastic process where the probability that a DNA molecule is amplified in a given cycle is less than one. This stochasticity may cause some sequences to have an artificially high RNA count. We note that assuming that the same reporter gene is used for each sample, the only difference in the sequence being amplified would be the barcode. Since barcodes are typically much shorter, it is unlikely to significantly alter the GC-content of the sequence, and therefore we do not discuss the effect of PCR sequence bias. Secondly, during RNA-Seq as well as the prior library preparation procedures such as RNA extraction and reverse transcription, we cannot ensure that every mRNA is extracted, converted to cDNA, and sequenced. Instead, in these steps, only a random subset of the original pool of mRNAs is sampled and included in the final sequencing dataset. As a result, a sequence may have an artificially low RNA count because some copies of the mRNA associated with that sequence are not sampled in one of the experimental steps.

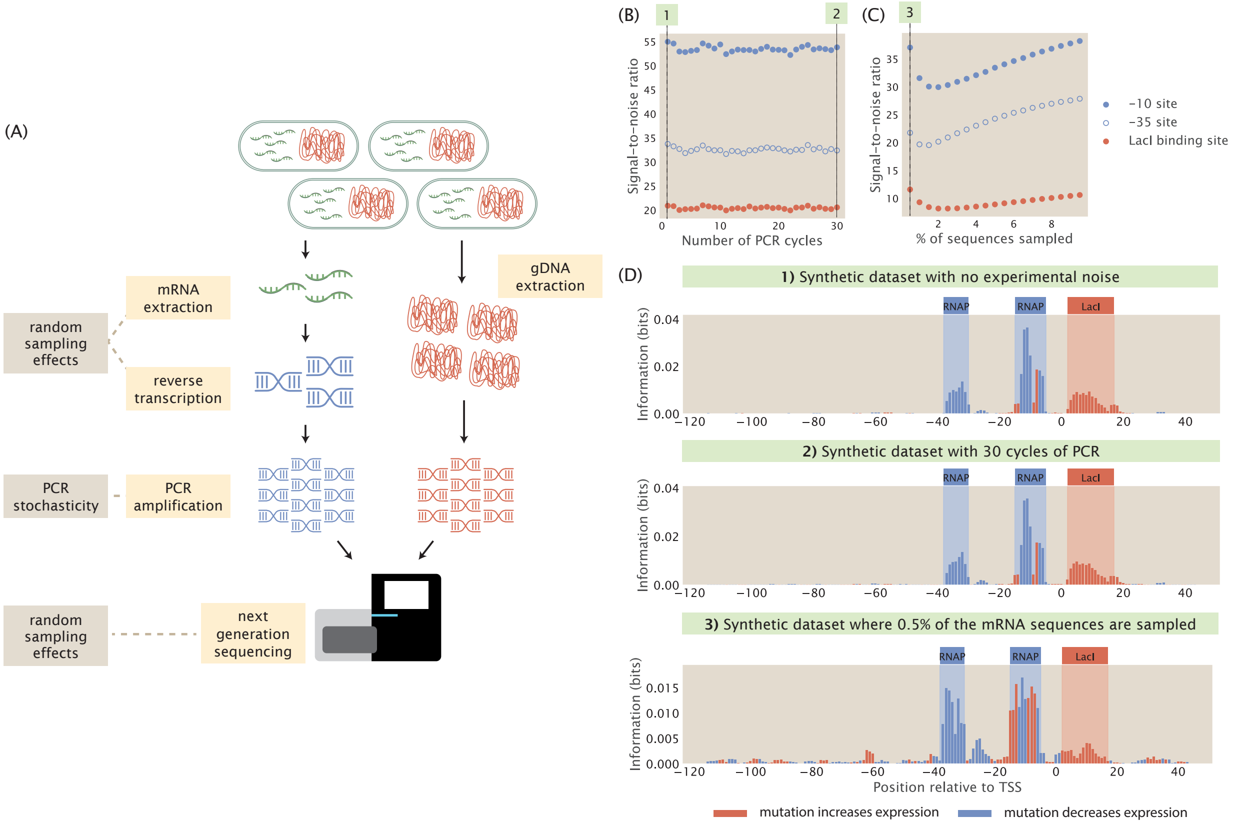


**Fig S10. Noise from experimental procedures in the Reg-Seq pipeline.** (A) The two main sources of noise in the experimental MPRA pipeline are stochasticity from PCR amplification and random sampling effects from RNA extraction, reverse transcription, and RNA-Seq. (B) Signal-to-noise ratio in the information footprints remains high when the number of PCR amplification cycles is increased. Here, $P_{\text{amp}} = 0.5$. Each data point is the mean of average mutual information across 20 synthetic datasets with the corresponding number of PCR cycles. The numbered labels correspond to footprints in (D). (C) Signal-to-noise ratio remains high when only a small percentage of the sequences are randomly sampled. Each data point is the mean of average mutual information across 20 synthetic datasets with the corresponding percentage of sampled sequences. The numbered labels correspond to footprints in (D). (D) Representative information footprints with no experimental noise, PCR stochasticity after 30 cycles, and random sampling effects after 0.5% of the RNA sequences are sampled.

We simulate these two sources of experimental noise in our computational pipeline. To simulate PCR with $n$ cycles of amplification, we start with the original mRNA counts predicted based on the probability of RNAP being bound. Subsequently, we model the number of sequences that are successfully amplified during each cycle using a Binomial distribution [60]. Hence, for each sequence variant,

$$n(j+1) = n(j) + B(n(j), P_{\text{amp}}), \tag{S43}$$

where $n(j)$ is the number of sequences of the promoter variant in cycle $j$, $B(n, P)$ models the Binomial distribution, and $P_{\text{amp}}$ is the probability that a sequence is successfully amplified in a cycle. We applied Eq S43 to calculate the final count of each sequence variant in a library. As shown in Fig S10(B) and S10(D), even when the probability of amplification is set to a low number of $P_{\text{amp}} = 0.5$, increasing the number of PCR cycles does not reduce the signal-to-noise ratio in information footprints. Therefore, we conclude that stochasticity in PCR does not contribute to significant levels of noise in information footprints.

To simulate the random sampling effect during RNA extraction, reverse transcription, and sequencing, we randomly draw a subset of promoter variants in the mutant library and we only consider the expression levels of the selected promoter variants when we calculate mutual information to build the information footprint. As shown in Fig S10(C) and S10(D), the levels of noise only becomes significant when less than 1% of the original pool of sequences is sampled. Therefore, random sampling effects are not a significant source of noise in information footprints either.

### S12 Appendix Modelling extrinsic noise using a Log-Normal distribution

In order to account for extrinsic noise, we choose to use a Log-Normal distribution to describe the copy number of RNAPs and repressors. Let $X$ be the copy number of the RNAP or the transcription factor, we define the Log-Normal distribution as

$$\log X \sim \text{Normal}\left(\log \mu, (\alpha \log \mu)^2\right). \tag{S44}$$

Since the goal of our analysis is to better formulate regulatory hypotheses based on real world data, we focus on levels of copy number fluctuations that are physiologically relevant. Assuming that the extrinsic noise in copy numbers primarily comes from asymmetrical partitioning during cell division, one method to measure fluctuations in transcription factor copy numbers between cells is the dilution method developed by Rosenfeld, Young et al. [61]. Based on Brewster et al. [62] who utilized the dilution method, transcription factor copy numbers typically vary by less than 20% of the mean copy number. Consider the proteomic measurements from Schmidt et al. [56] and Balakrishnan et al. [63], the coefficient of variation for transcription factor copy numbers is less than 2 even across very different growth conditions. With these empirical data, we can then define our distributions of copy numbers to respect the known levels of fluctuation. Given the known mean and variance of a Log-Normal distribution, we can derive that the coefficient of variation is given by

$$\text{CoV}(X) = \frac{\mathbb{E}(X)}{\text{Var}X} = \sqrt{e^{(\alpha \log \mu)^2} - 1}.$$

Rearranging this expression, we can write down $\alpha$ in terms of $\mu$ and $\text{CoV}(X)$, where

$$\alpha = \frac{\sqrt{\log\left[\text{CoV}(X)^2 + 1\right]}}{\log \mu}$$

This means that we can derive a Log-Normal distribution that obeys the empirical mean and coefficient of variation for transcription factors. In Fig 12, we simulated noisy synthetic datasets using Log-Normal distributions with $P = 5000$ and $R = 100$ as the mean copy numbers and a range of coefficients of variation from 0.1 to $10^2$, which covers the levels of fluctuations that are physiologically relevant. In Fig S11, we show the distributions of copy numbers given the three levels of fluctuations that we particularly discuss in Sec 3.1. Note that when the coefficient of variation is set to 100, the copy number of RNAPs can reach as high as $10^9$ and the copy numbers of repressors can reach as high as $10^7$, both of which are very unrealistic levels of copy numbers.

### S13 Appendix Extrinsic noise under low signal

In Sec 3.1, we explored the effect of extrinsic noise while keeping all other parameter at their standard values. That is to say, we expect that the fluctuations in copy numbers should not drastically change the level of signal in the footprints. However, one reasonable suspicion is that when the signal from binding events is sufficiently low, even low levels of noise may affect our interpretation of information footprints and expression shift matrices. To examine whether this is the case, we built footprints with lowered binding energy for the repressor while allowing copy numbers to fluctuate. As shown in Fig S12, even when the signal is low and the coefficient of variation is set to as high as 100, we can still identify signal at the repressor binding site. Therefore, it seems that this is not an issue unless the level of noise is unrealistically high.

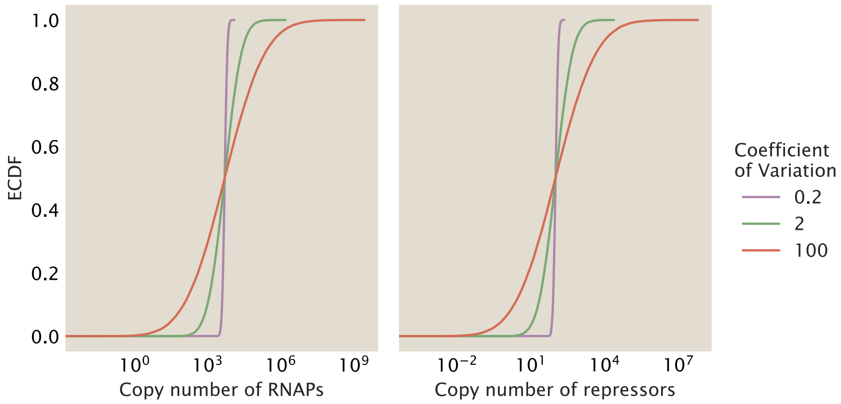


**Fig S11. Modelling the copy number of RNAPs and repressors using a Log-Normal distribution.** CDFs for the copy numbers of RNAP and repressors modelled using a Log-Normal distribution and under three different levels of coefficient of variation.

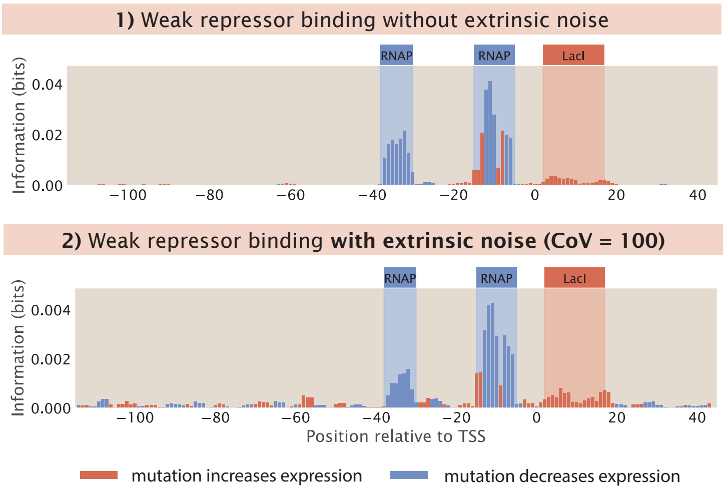


**Fig S12. Effects of extrinsic noise on information footprints with weak repressor binding.** For both footprints, the repressor binding energy $\Delta\varepsilon_{\mathrm{rd}} = -11\, k_B T$. In the top information footprint, no extrinsic noise is introduced. In the bottom information footprint, the Log-Normal distribution from which RNAP and repressor copy numbers are drawn has a coefficient of variation of 100.

### S14 Appendix Extrinsic noise for different architectures

To see if our investigation of extrinsic noise is generalizable, we test the cases where there are copy number fluctuations under the other four common regulatory architectures (simple activation, double repression, double activation, and repression-activation). As shown in Fig S13, the signal-to-noise ratio remains high regardless of the architecture even when copy numbers are allowed to fluctuate to 10 times above or below the average copy number.

### S15 Appendix Statistical weights for the simple activation promoter based on spanning trees

To derive the statistical weights of all states using the graph-theoretic approach introduced in Sec 4, we make use of the Matrix Tree Theorem, which states that at steady state, the probability of state $i$ is proportional

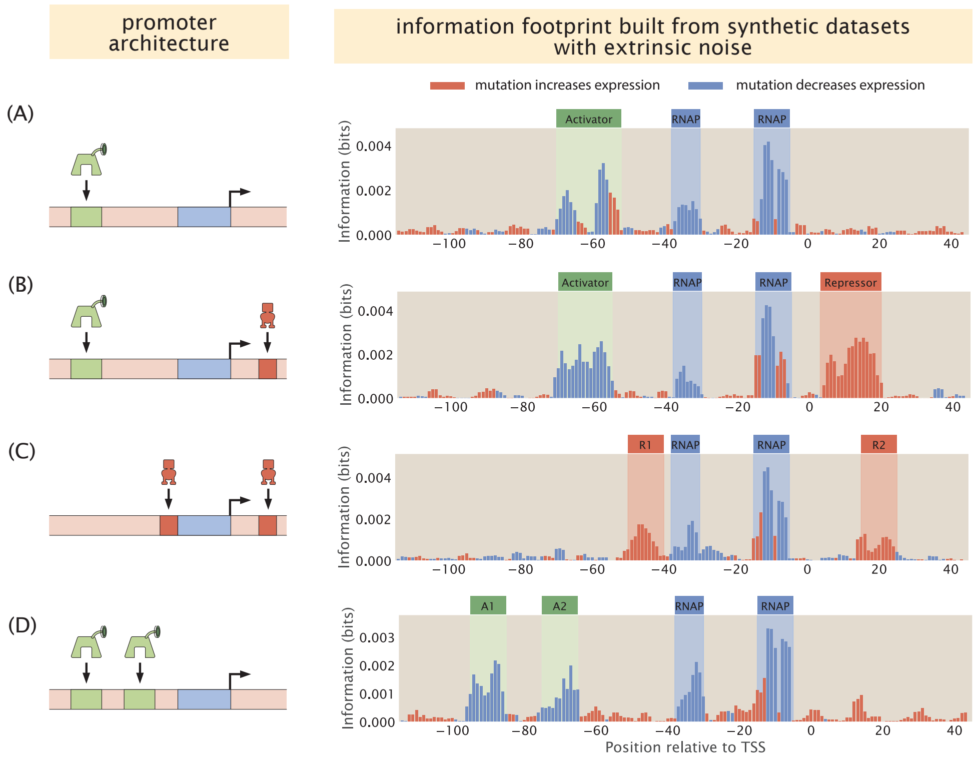


**Fig S13. Information footprints built from synthetic datasets with extrinsic noise under common regulatory architectures.** In all four plots, the coefficient of variation for RNAP and transcription copy numbers is set to 10. Panel (A) to (D) show the footprints from the simple activation architecture, repression-activation architecture, double repression architecture, and the double activation architecture, respectively.

to the sum of products of rate constants across all spanning trees rooted in the vertex representing state $i$. By definition, a spanning tree rooted in vertex $i$ is a subgraph that (1) contains all vertices in the original graph and (2) has all edges incoming in vertex $i$. The graph describing a simple activation promoter is shown in Fig 15. As shown in Fig S14, we can enumerate all spanning trees for each of the four vertices. This gives us the following statistical weights for each of the promoter states

$$\rho_{\mathrm{E}} = k_{\mathrm{AE}}\, k_{\mathrm{PE}}\, k_{\mathrm{AP,P}} + k_{\mathrm{A,AP}}[\mathrm{P}]\, k_{\mathrm{AP,P}}\, k_{\mathrm{PE}} + k_{\mathrm{PE}}\, k_{\mathrm{AP,P}}\, k_{\mathrm{AE}} + k_{\mathrm{P,AP}}[\mathrm{A}]\, k_{\mathrm{AP,P}}\, k_{\mathrm{AE}} \tag{S45}$$

$$\rho_{\mathrm{P}} = k_{\mathrm{AE}}\, k_{\mathrm{EP}}[\mathrm{P}]\, k_{\mathrm{AP,P}} + k_{\mathrm{EP}}[\mathrm{P}]\, k_{\mathrm{A,AP}}[\mathrm{P}]\, k_{\mathrm{AP,P}} + k_{\mathrm{AP,A}}\, k_{\mathrm{AE}}\, k_{\mathrm{EP}}[\mathrm{P}] + k_{\mathrm{EA}}\,[\mathrm{A}]\, k_{\mathrm{A,AP}}[\mathrm{A}]\, k_{\mathrm{AP,P}} \tag{S46}$$

$$\rho_{\mathrm{A}} = k_{\mathrm{AP,P}}\, k_{\mathrm{PE}}\, k_{\mathrm{EA}}[\mathrm{A}] + k_{\mathrm{EP}}[\mathrm{P}]\, k_{\mathrm{P,AP}}[\mathrm{A}]\, k_{\mathrm{AP,A}} + k_{\mathrm{PE}}\, k_{\mathrm{EA}}[\mathrm{A}]\, k_{\mathrm{AP,A}} + k_{\mathrm{P,AP}}[\mathrm{A}]\, k_{\mathrm{AP,A}}\, k_{\mathrm{EA}}[\mathrm{A}] \tag{S47}$$

$$\rho_{\mathrm{AP}} = k_{\mathrm{AE}}\, k_{\mathrm{EP}}[\mathrm{P}]\, k_{\mathrm{P,AP}}[\mathrm{A}] + k_{\mathrm{EP}}[\mathrm{P}]\, k_{\mathrm{P,AP}}[\mathrm{A}]\, k_{\mathrm{A,AP}}[\mathrm{P}] + k_{\mathrm{PE}}\, k_{\mathrm{EA}}[\mathrm{A}]\, k_{\mathrm{A,AP}}[\mathrm{P}] + k_{\mathrm{EA}}[\mathrm{A}]\, k_{\mathrm{A,AP}}[\mathrm{P}]\, k_{\mathrm{P,AP}}[\mathrm{A}] \tag{S48}$$

Finally, we can calculate the probability of each state by taking the weight of each state and dividing by the sum of all weights

$$p_{\mathrm{E}} = \frac{\rho_{\mathrm{E}}}{\rho_{\mathrm{E}} + \rho_{\mathrm{P}} + \rho_{\mathrm{A}} + \rho_{\mathrm{AP}}} \tag{S49}$$

$$p_{\mathrm{P}} = \frac{\rho_{\mathrm{P}}}{\rho_{\mathrm{E}} + \rho_{\mathrm{P}} + \rho_{\mathrm{A}} + \rho_{\mathrm{AP}}} \tag{S50}$$

$$p_{\mathrm{A}} = \frac{\rho_{\mathrm{A}}}{\rho_{\mathrm{E}} + \rho_{\mathrm{P}} + \rho_{\mathrm{A}} + \rho_{\mathrm{AP}}} \tag{S51}$$

$$p_{\mathrm{AP}} = \frac{\rho_{\mathrm{AP}}}{\rho_{\mathrm{E}} + \rho_{\mathrm{P}} + \rho_{\mathrm{A}} + \rho_{\mathrm{AP}}} \tag{S52}$$

In particular, since $A$ and $AP$ are the transcriptionally active states, the probability that the promoter is on 1451
is given by $p_{\text{active}} = p_{\text{A}} + p_{\text{AP}}$. 1452

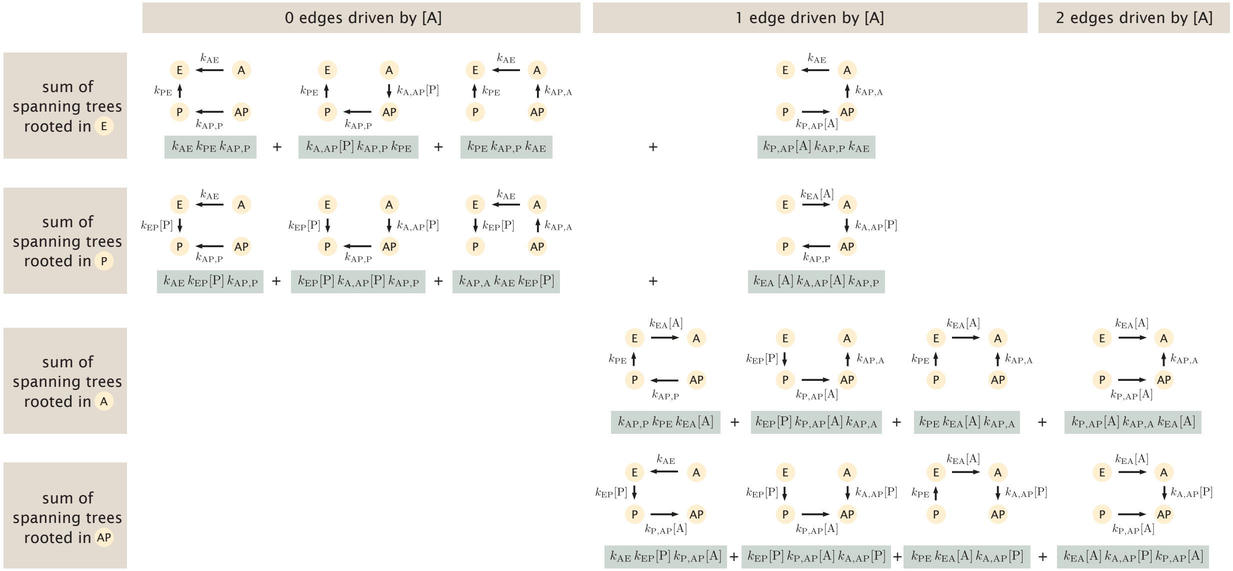


**Fig S14. Deriving statistical weights of promoter states using spanning trees.** Each row corresponds to a different root for the spanning trees. The columns are grouped based on the number of edges in the spanning tree that depend upon the concentration of the activator. The figure is adapted from Mahdavi, Salmon et al. [4].
